# Convolutional Neural Networks & Neuroscience: A Tutorial Introduction for The Rest of Us

**DOI:** 10.64898/2026.03.09.710521

**Authors:** Matteo De Matola, Giorgio Arcara

**Affiliations:** Center for Mind/Brain Sciences— CIMeC, University of Trento, Rovereto (TN), 38068, Italy; Department of General Psychology, University of Padova, Padova, 35131, Italy; IRCCS San Camillo Hospital, 30126 Venice, Italy

## Abstract

Convolutional neural networks (CNNs) are a class of artificial neural networks (ANNs). Since the early 2010s, they have been widely adopted as models of primate vision and classifiers of neuroimaging data, becoming relevant for a wealth of neuroscientific fields. However, the majority of neuroscience researchers come from soft-science backgrounds (like medicine, biology, or psychology) and do not have enough quantitative skills to understand the inner workings of A/CNNs. To avoid undesirable black boxes, neuroscientists should acquire some rudiments of computational neuroscience and machine learning. However, most researchers do not have the time nor the resources to make big learning investments, and self-study materials are hardly tailored to people with little mathematical background. This paper aims to fill this gap by providing a concise but accurate introduction to CNNs and their use in neuroscience — using the minimum required mathematics, neuroscientific analogies, and Python code examples. A [companion Jupyter Notebook](https://github.com/matteo-d-m/cnns-tutorial) guides readers through code examples, translating theory into practice and providing visual outputs. The paper is organised in three sections: The Concepts, The Implementation, and The Biological Plausibility of A/CNNs. The three sections are largely independent, so readers can either go through the entire paper or select a section of interest.

## 1 Introduction

Artificial neural networks (ANNs) are mathematical models of networks of interacting biological neurons [1–4]. Over the last ten years, their implementations in digital computers have proven able to simulate some aspects of human cognition, such as vision [5–7], language [8–11], learning [12], and decision-making [13–16]. Such results originated mostly from computer science departments and technology companies, but inevitably attracted the interest of the neuroscience community — with more than 25000 papers of neuroscientific interest published over the last five years ^1^. The increasing popularity of ANNs entails a need to understand their inner workings, but the average neuroscientist has little or no training in mathematics and computer science. The literature and the Internet provide a wealth of excellent training resources, in both traditional formats like books and more innovative formats like online courses (for example, the [Neuromatch Academy](https://neuromatch.io/)). Unfortunately, the fast pace and multiple commitments of academic careers leave little opportunities for continuous learning, and studying a book or following a course might require a disproportionate investment for researchers that seek an introduction to the topic but cannot afford to become experts. Moreover, learning resources tend to be either mathematically sophisticated (therefore, not accessible to people with a poor quantitative background) or excessively simplistic (therefore, less insightful and formally correct than they could be). More often than not, knowledge is scattered across different places and theoretical resources are separated from technical resources, forcing practitioners to do two separate learning efforts: one to find, select and learn *theoretical* knowledge, and another to find, select and learn *practical* knowledge. In parallel, the availability of user-friendly software tools like PyTorch [17] and pre-trained models like AlexNet [5] allows practitioners to use complex ANNs without having to know their inner workings — which is desirable from the point of view of productivity, but certainly undesirable from the point of view of scientific rigour. Overall, this situation calls for a reform of neuroscience training programs — a topic that has been treated extensively elsewhere [18–21]. In the meantime, learning materials that strike the balance between formal rigour, usability, and suitability to a non-quantitative audience might help on the short term. This paper aims to provide a simple yet rigorous introduction to ANNs, using a combination of text, simple mathematical formulas, images, and Python code. The authors are neuroscience researchers that struggled to become conscious users of advanced data analysis methods, and the target audience are fellow researchers (or students) walking the same path. The only prerequisites for this paper are very basic mathematics (essentially, sums and products) and the ability to read (but not necessarily write) computer code in a high-level, user-friendly programming language like Python. While the paper’s goal is to be a self-contained learning resource, some concepts (for example, matrices) are necessarily given only an essential treatment. These concepts will be highlighted (e.g., <u>matrix</u>) as a reminder that they might require further study. However, readers should be able to get a full grasp of the paper without delving deeper into any specific concept.

After introducing ANNs in general, the paper will focus on convolutional neural networks (CNNs): a class of ANNs that are widely used as models of neurocognitive functions (in particular, visual perception in the primate ventral visual stream [22]) and as tools for neuroimaging data analyses (for example, segmenting and classifying magnetic resonance images [23–25] or electroencephalography time windows [26–29]). To close the gap between theory and practice, the main text of this paper is complemented by Python code blocks that implement the theory. Moreover, the second section of the paper is entirely dedicated to introducing and explaining PyTorch, the go-to Python library for ANN practitioners. All the code included in this paper (and much more) can be run in the associated Jupyter Notebook^2^, which is publicly available via GitHub at this link under a permissive license.

## 2 The Concepts

### 2.1 Core Concept One: Artificial Neural Networks

As it happens with most things, there is no single way to conceptualise ANNs. In fact, there are at least three ways to think about them: a mathematical way, a computational way, and a neuroscientific way. The mathematical way conceptualises ANNs as combinations of <u>multivariate parametric</u> linear and non-linear functions which are optimised by some variation of <u>stochastic gradient descent</u>^3^. The computational way conceptualises ANNs as computer programs that take in inputs, transform them according to some rules, and return outputs. Finally, the neuroscientific way treats ANNs as models of cognitive functions in the primate brain, using neuroanatomical and neurophysiological jargon to give them meaning. All three ways are valid, and the choice between them depends entirely on the use case or the target audience. This paper is aimed at neuroscientists, so it emphasises the neuroscientific way. However, it uses mathematical tools and concepts when needed to avoid black boxes.

#### 2.1.1 The Artificial Neuron

The artificial neuron is the cornerstone of artificial neural networks. It is a mathematical model (basically, one equation) that represents a neuron as a <u>weighted sum</u>^4^ of its inputs. Given:

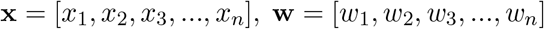

calculate:

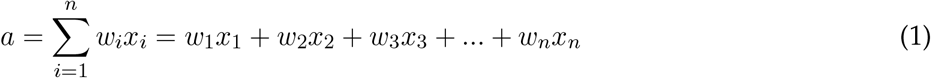

where **x** is a <u>vector</u> that stores *n* input values, **w** is a vector that stores *n* corresponding weights, and *a* is the neuron’s activation. For the purpose of this paper, a vector can be defined as a container of numbers that represent an object: for example, a vector of two coordinates *x, y* represents a point on a plane, while a vector of *n* values *x*_1_, *x*_2_, *x*_3_, …, *x*_*n*_ can represent the luminosity of *n* dots.

The notation 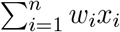 means that all the values of **W** numbered from *w*_1_ to *w*_*n*_ (that is, all the *n* components of vector **w**) are multiplied with the corresponding values of *x*, then the results of those multiplications are summed (Σ).

Equation 1 is meant to model three biological facts:

- Vector **x** models the fact that real neurons receive inputs from multiple synapses. If there was a single synapse, there would not be a vector but a single number
- Vector **w** models the fact that different synapses have different strengths, meaning that the corresponding inputs have different weights (i.e., importance, salience, relevance, or any other way of calling it). If all synapses had the same strength, **w** would contain multiple copies of the same number
- *a* models the fact that postsynaptic potentials (and *action* potentials) are a sum of the local potentials generated by synaptic activity

An artificial neuron may also have a *bias* term (*b*) to represent its baseline activity:

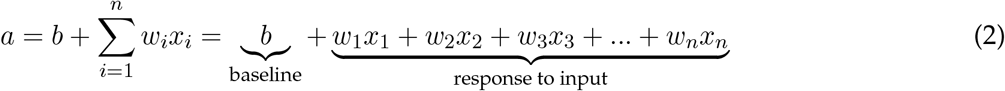

Code Example 1 shows a Python implementation of an artificial neuron that takes in a random threedimensional input. Note that in linear algebra jargon, the weighted sum of inputs is the *dot product* between vector **w** and vector **x**. Therefore, the corresponding NumPy [30]^5^ function is called ‘dot’.

##### Code Example 1: The Artificial Neuron

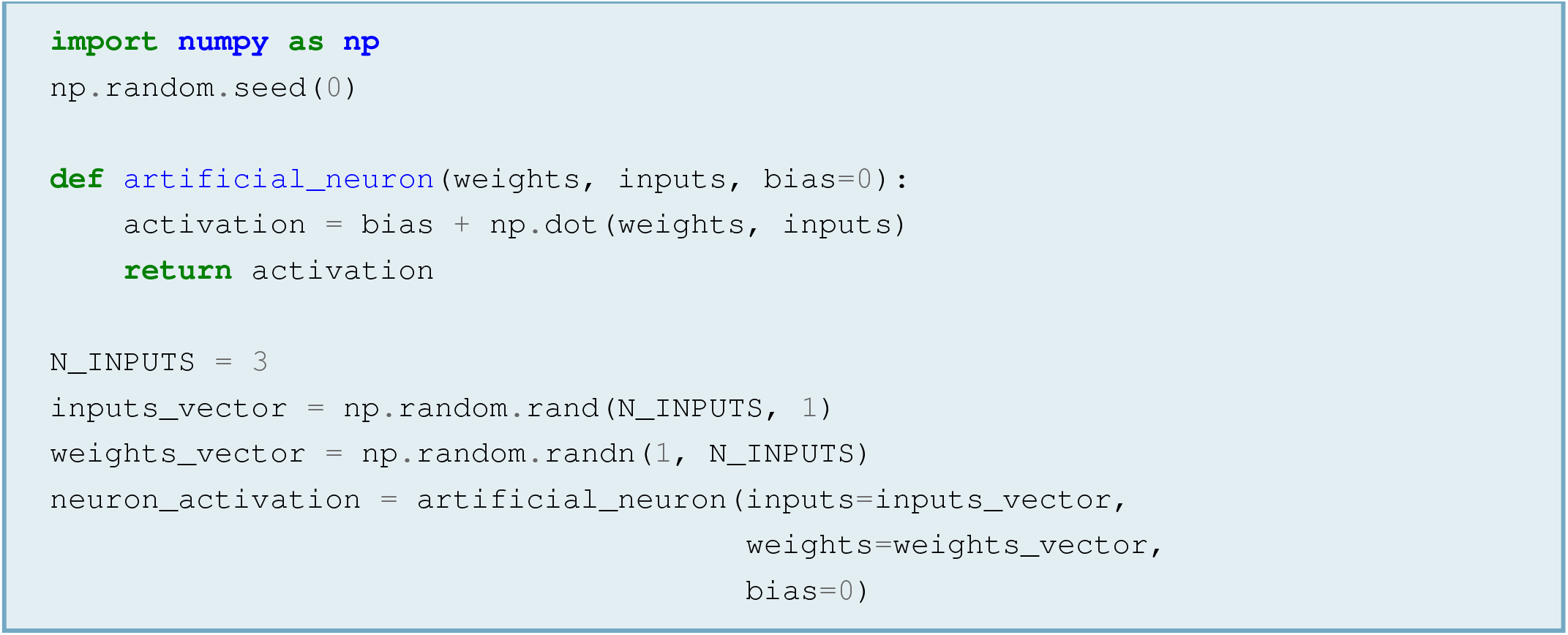

#### 2.1.2 The Activation Function

The computation of a neuron’s activation is usually followed by a non-linear function called the *activation function* (*f*):

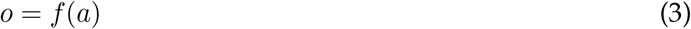

where *a* is the neuron activation (i.e., the weighted sum of its inputs), *f* is some non-linear function and *o* is the final output of the neuron. The role of the activation function is to account (or at least, try to account) for the non-linear nature of true neuronal activity [31, 32] The ANNs literature contains a wealth of activation functions, of which a classic example is the sigmoid (or *logistic*) function:

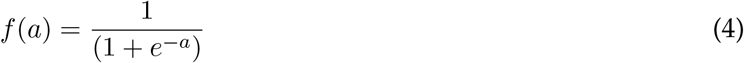

While it might look complex, the sigmoid does little more than compressing the input between 0 and 1, producing an output that can be interpreted in terms of probability (for example, probability that a neuron fires in response to a given input). The sigmoid is extremely simple to implement in Python, as shown in Code Example 2.

##### Code Example 2: Sigmoid Function

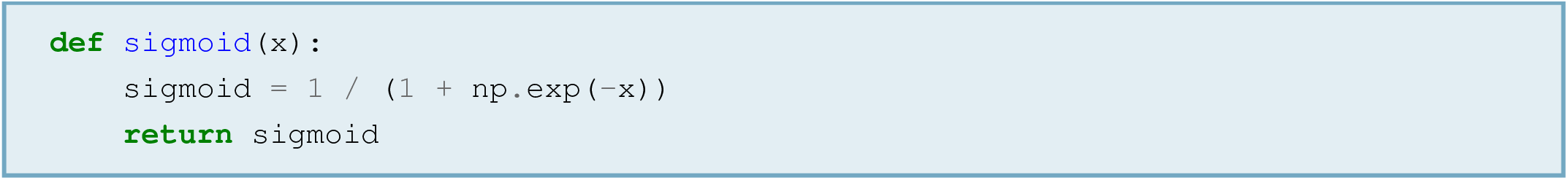

The sigmoid has been important in the past, but is not much used nowadays. A more modern activation function is the rectified linear unit (ReLU), which sets all negative values to zero and leaves non-negative values as they were (Figure 1). In practice, this means that ReLU checks the value of *x* and compares it to zero to see if it is positive or negative. If *x* is negative, 0is the largest number and will be ReLU’s output; while if *x* is positive, *x* is the largest number and will be ReLU’s output. This can be written as:

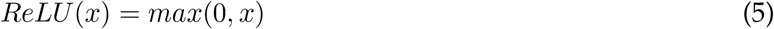

**Figure 1.**
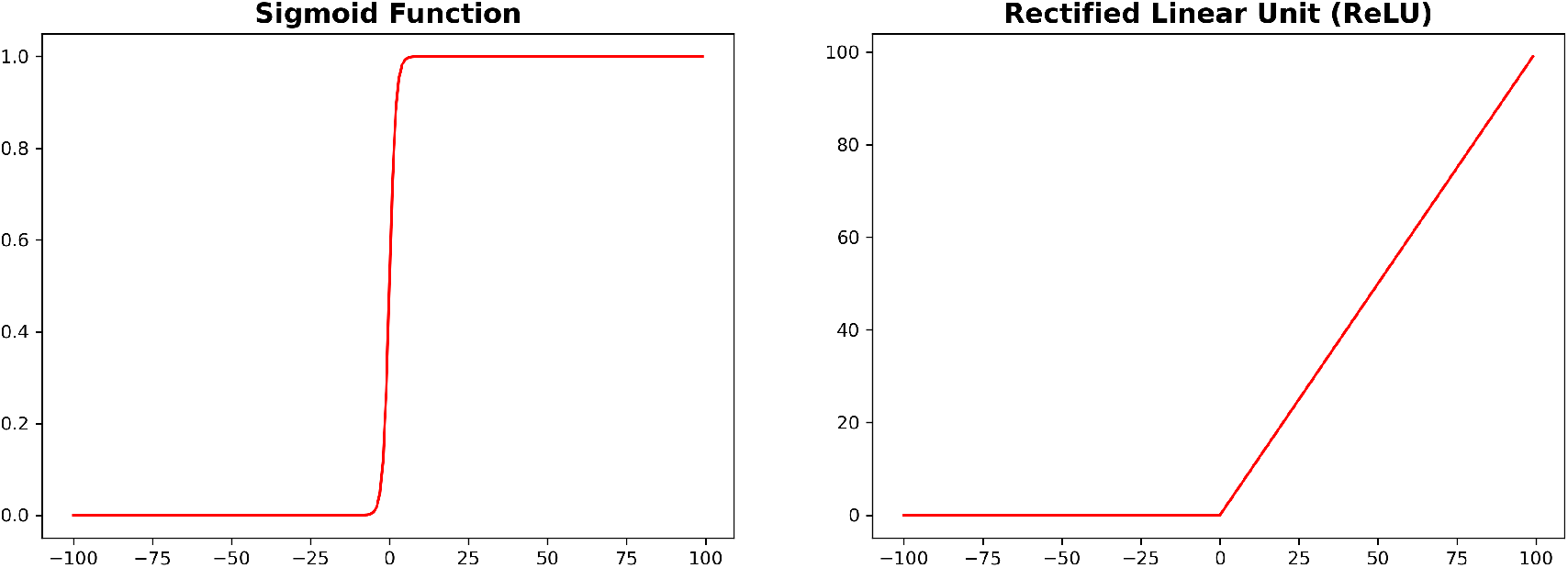
A sigmoid (left) and a ReLU (right) function

This compact formulation is adopted in Code Example 3:

##### Code Example 3: Rectified Linear Unit (ReLU)

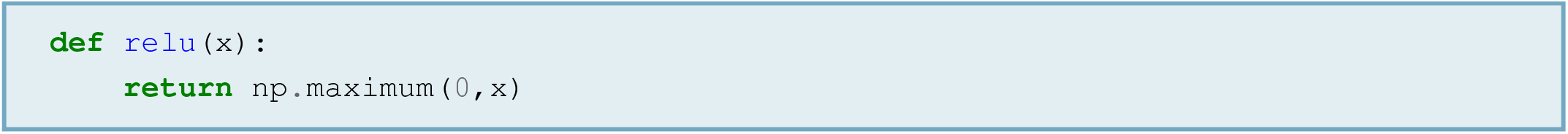

#### 2.1.3 From Neuron to Network

The previous section showed that a biological neuron can be very simply modelled as the dot product (that is, a weighted sum) between two vectors — **x** and **w** — which represent a set of inputs and a set of corresponding weights. To model a larger set of interacting neurons (that is, a *network*), the vector of inputs (**x**) must be multiplied by a whole *matrix* of weights (**W**). This matrix will have one row for each neuron and one column for each input to that neuron. Given:

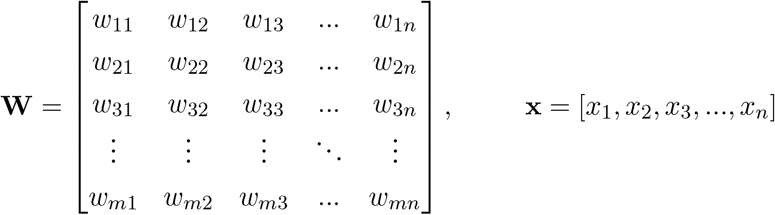

calculate:

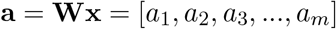

where **x** is a vector that stores *n* input values, **W** is a matrix that stores *n* corresponding weights (over columns) for each of *m* neurons (over rows), and **a** is a vector that stores one activation value for each of *m* neurons.

Each element of **a** is the dot product between **x** and one row of the weights matrix, as per the equation below:

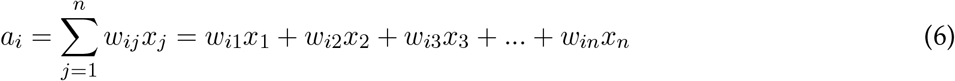

where *a*_*i*_ is the activation value of the *i*^*th*^ neuron. Code Example 4 shows the Python implementation of a very simple neural network with three neurons and a three-dimensional random input. As can be seen, the implementation is the exact same as the artificial neuron’s, with the only difference that the weights are now a matrix of shape `neurons x inputs`.

##### Code Example 4: Artificial Neural Network (ANN)

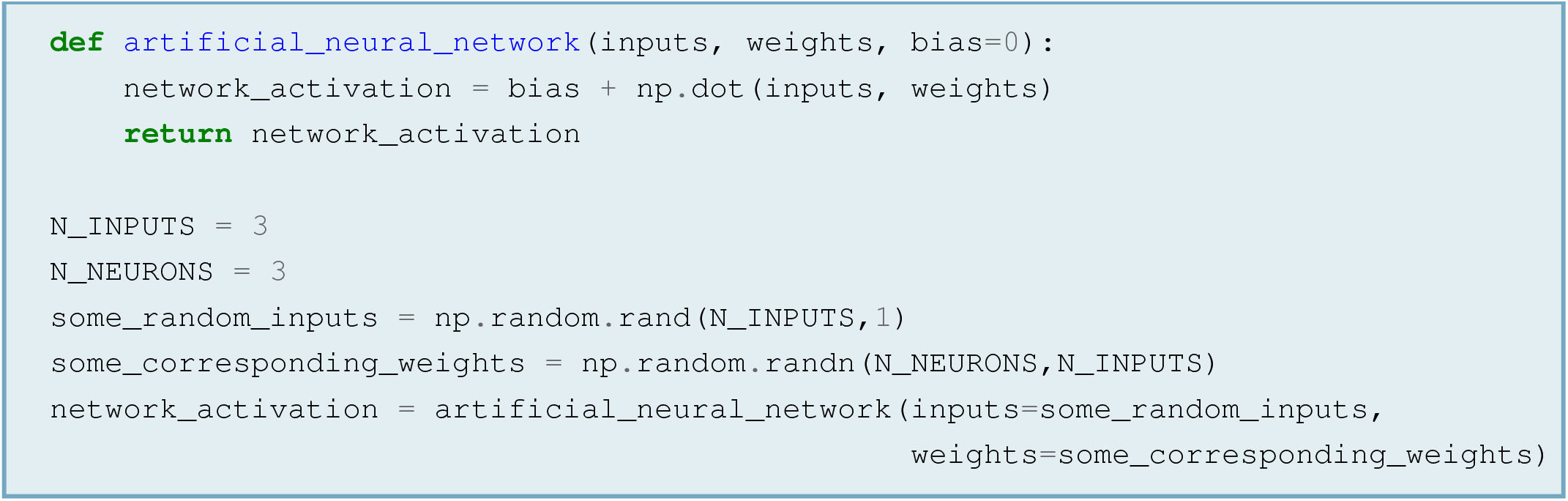

#### 2.1.4 From Network to Deep Network

The multiplication between a matrix of coefficients and a a vector of inputs yields the activity of a set of neurons — which, in neural networks jargon, is called a *network layer*. A network layer is analogous to a cortical area, where multiple neurons respond to incoming signals through a set of synapses with different weights. However, this is not enough to represent a situation where *multiple* sets of neurons communicate with each other, as is the case when a brain area sends a signal to another one through multiple intermediate relay stations. Neural networks simulate this situation by *concatenating* multiple layers, such that the output of a set of neurons becomes the input to another one and eventually, the output of the whole network is shaped by the activity of each intermediate layer. Code Example 5 shows a simple example of a three-layers neural network that takes in a three-dimensional vector of random data and processes it with three layers of three neurons each. The activation function is a ReLU, but it could be anything:

##### Code Example 5: Deep Artificial Neural Network (dANN)

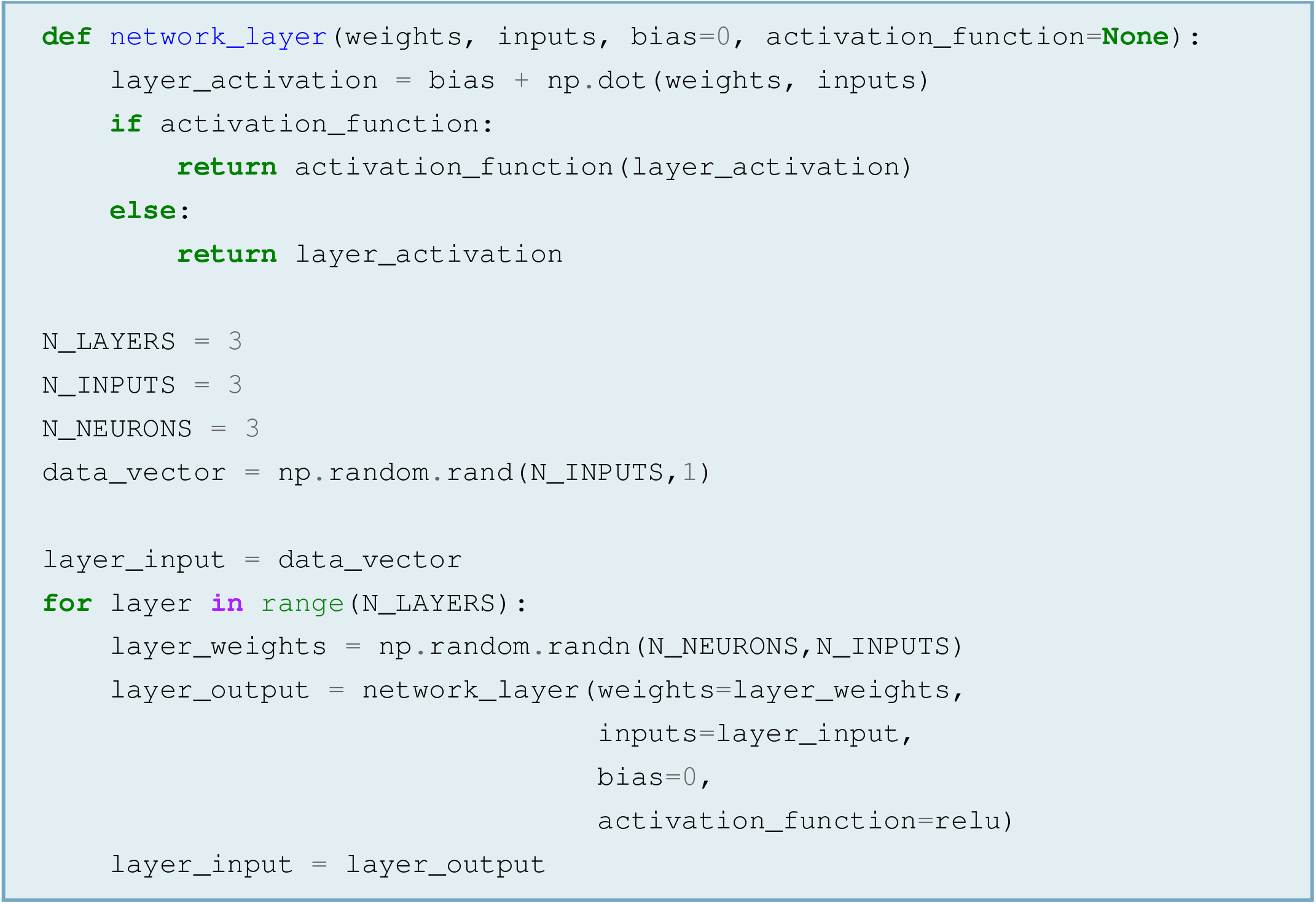

Deep neural networks are often introduced to learners through a graphical representation where each artificial neuron is a circle and each interaction between them is an arrow (Figure 2). While they might seem very abstract, such graphics represent the very multiplication between a matrix of weights and a vector of data (or neuronal activations). The stacks of circles represent the vectors, and the arrows represent the effect of multiplication with weight matrices. At each layer in the network (that is, each stack of circles), each neuron (circle) sends one arrow to each neuron in the next layer, in such a way that all neurons in one layer send information to all neurons in the next.

**Figure 2.**
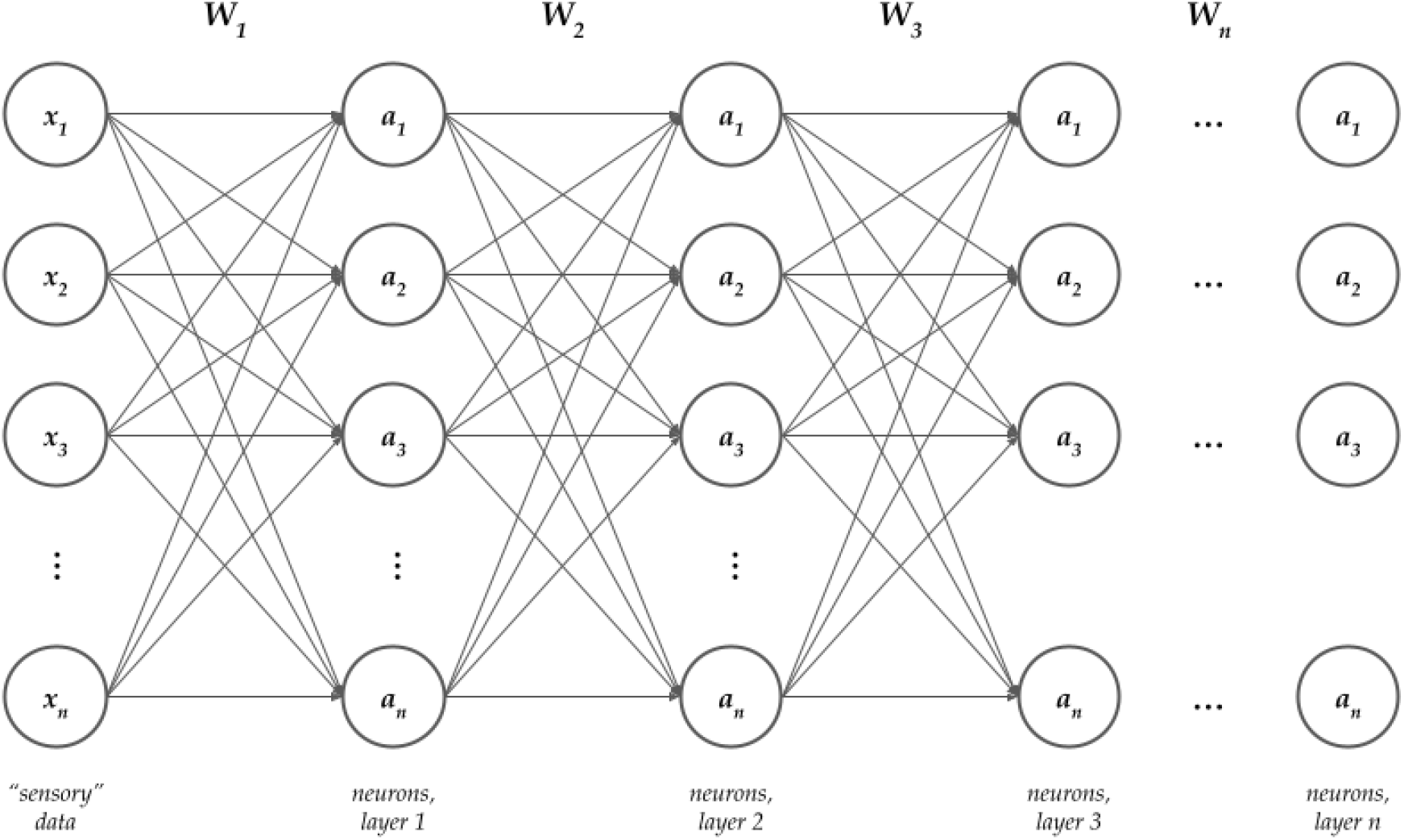
Neural network diagrams represent matrix-vector multiplications

In the equation below, vector **x** represents data (say, luminosity values in a tiny image with three pixels) and vector **a** represents neuronal activations (for example, of three retinal cells). The three colored dots next to the components of **a** emphasise that all components of **x** are involved in generating all components of **a**:

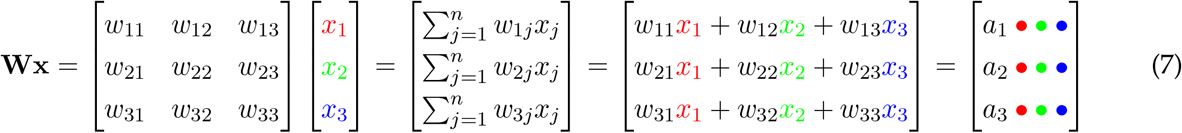

Analogously, all the neurons in any layer of Figure 2 send arrows to all the neurons in the next layer:

#### 2.1.5 The Problem of Bidimensional Inputs

Standard network layers work well for vector inputs of the form **x** = [*x*_1_, *x*_2_, *x*_3_, …, *x*_*n*_] — which, in practice, might be EEG data from a given electrode (one number per time point):

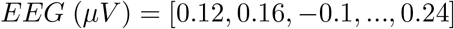

or the mean rent in a Rovereto city district at any given time (one number per district):

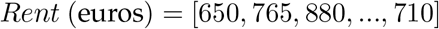

or anything else that can be described by a one-dimensional array. But, what happens with two-dimensional inputs? For example, what happens with EEG data from several electrodes, where each electrode is one row of a matrix?

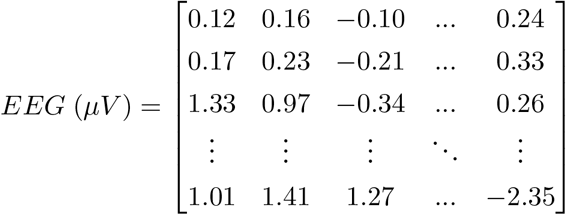

In principle, bidimensional inputs could be multiplied by a weights matrix like their vector counterparts. Given:

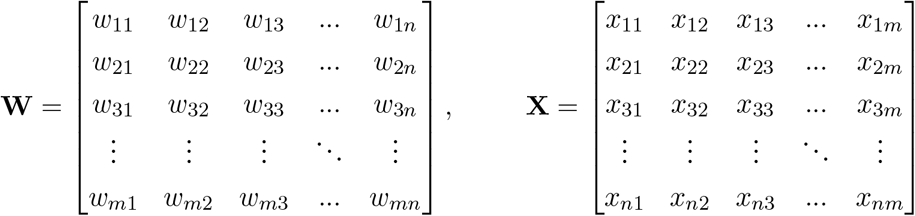

we could calculate:

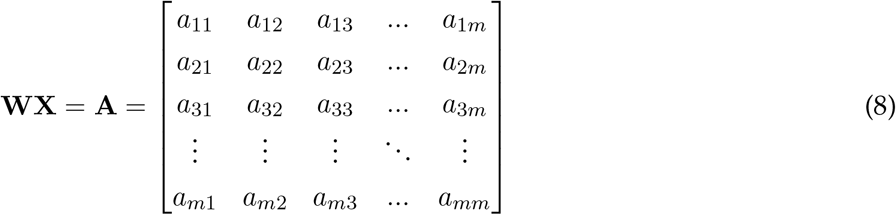

where **X** is a matrix that stores *n* · *m* intensity values of a bidimensional input, **W** is a matrix that stores *m* · *n* corresponding weights, and **A** is a matrix that stores *m* · *m* activation values for as many neurons. Here, each element of **A** would be the dot product between one row of **W** and one column of **X**:

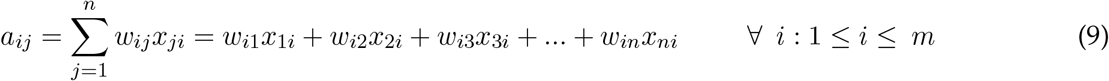

where *a*_*ij*_ would be the activation of the neuron found in position *i, j* within the bidimensional arrangement of neurons represented by **A**. This neuron would respond to the *j*^*th*^ column of the input matrix through the weights found in the *i*^*th*^ row of the weights matrix. Clearly, this strategy has no straightforward neuroscientific interpretation, as the algebraic machinery involved in **A** = **WX** does not seem to represent any major neurophysiological process. Moreover, this machinery does nothing to emphasize the *compositional structure* that data may have. The next section will explain what this means.

### 2.2 Core Concept Two: Convolutional Neural Networks

Grid-like inputs — that is, inputs arranged as lines, squares, or cubes — are often a composition of local structures, or *features*: patterns that appear in given parts of the grid and are its fundamental building blocks. In a one-dimensional grid like a single EEG channel, features can be bursts of activity circumscribed to specific time windows. In a two-dimensional grid like a multi-channel EEG, features can be a distribution of such patterns over several neighbouring channels. In an image, features can be geometric primitives like edges and circles. Whatever the nature of the features, a neural network can extract them from input data with a *convolution*. In simple terms, a convolution is an operation that involves five steps:

1. Take a weights matrix that is smaller than the input (**W**_**small**_). This is usually referred to as the convolution *kernel* or convolution *filter*
2. Overlay **W**_**small**_ to an equally small patch of the input
3. Multiply **W**_**small**_ and the corresponding input patch, element by element
4. Sum all the element-wise products. This will be a neuron’s activation
5. Repeat on a neighbouring patch to get another neuron’s activation
6. Continue until covering the whole input

Points 5 and 6 mean that the convolution kernel operates in a moving window fashion (cf. Figure 3).

**Figure 3.**
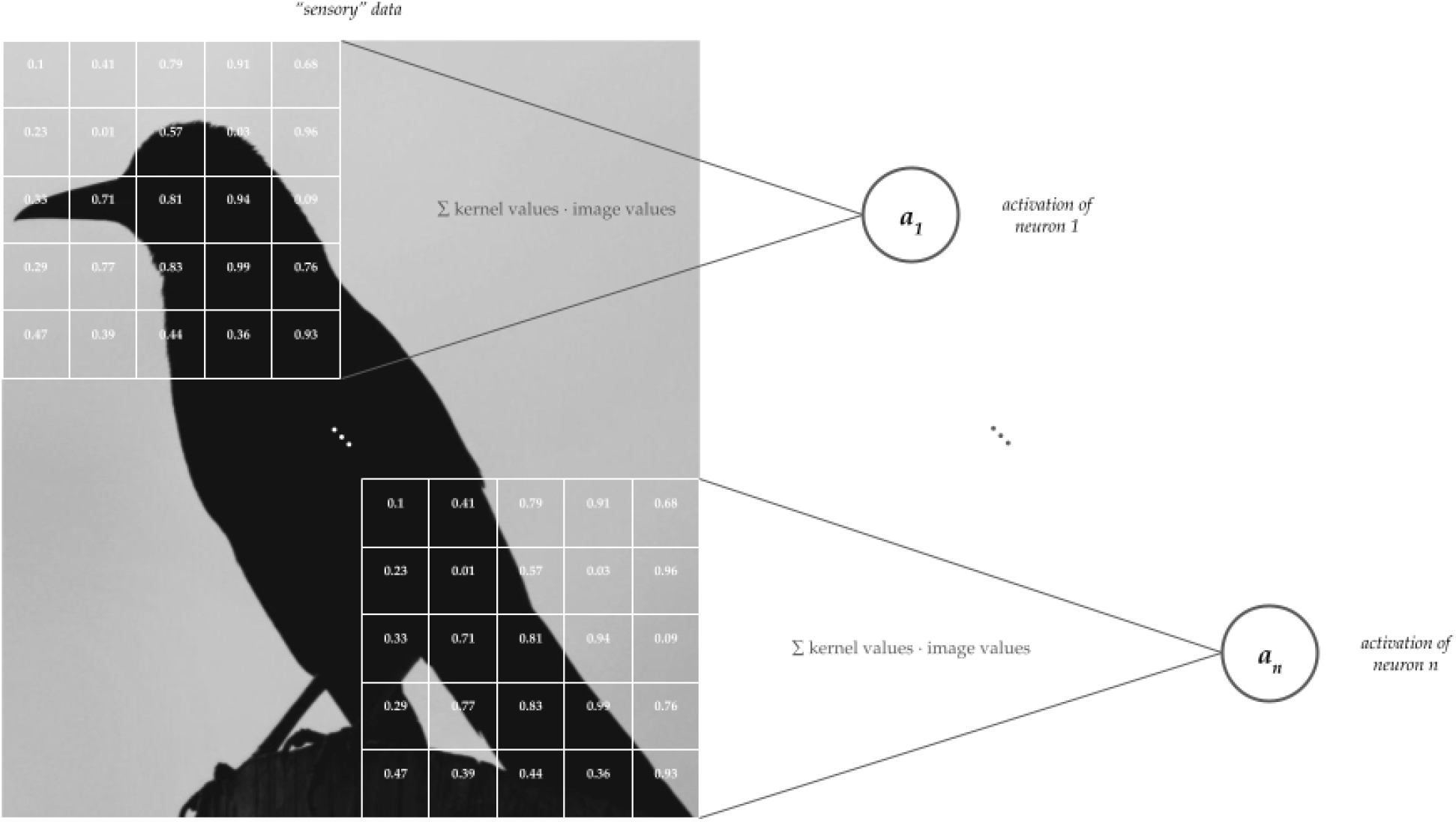
A convolution over a black-and-white image (bird photo by Joel Naren on Unsplash). The white matrix is the convolution kernel, which is multiplied with the underlying image patch element-wise. After multiplication, the element-wise products are summed and possibly, fed through an activation function, yielding the activation of one neuron. The kernel slides over the whole image and element-wise products are computed at each step. In this example, the top neuron gets activation *a*_1_ in response to the convolution with the top-left portion of the image, while the bottom neuron gets activation *a*_*n*_ in response to the convolution with the bottom-right portion of the image. Other neurons (and the corresponding convolution steps) are omitted to avoid overcrowding the figure.

There are three things about convolution that must be kept in mind:

1. The convolution of a grid-like input with some kernel is a special type of network layer that is called a *convolutional layer*. As it happens for standard network layers, the outputs of convolutional layers can be fed into arbitrary activation functions (for example, a sigmoid or a ReLU) and concatenated into a deep network (cf. 2.1.4)
2. From a strictly computational perspective, convolution is equivalent to taking the dot product between a patch of input and a set of weights. The only difference with a standard dot product is that the inputs and weights are topologically organised as a grid — that is, the spatial distribution of inputs and weights matters. The result of this topologically organised dot product can be seen as the activation of a sensory neuron, and the corresponding patch of input can be seen as the neuron’s receptive field (Figure 3)
3. Clearly, a convolution kernel returns one output for each of the *n* steps it takes over the input. Therefore, the final result of convolution is a grid of *n* activation values representing *n* neurons that respond to *n* different regions of the input. As these regions are supposed to contain *features*, the grids of activation values are called *feature maps*

The Jupyter Notebook associated to this paper ([GitHub link](https://github.com/matteo-d-m/cnns-tutorial)) includes Python code to implement a convolution using basic tools from NumPy, the go-to Python library for matrix operations. Understanding its content requires to know three terms: *kernel, stride*, and *padding*. As mentioned above, the *kernel* is merely a weights matrix. Like its synonym *filter*, the term *kernel* is inherited from the field of signal processing, where convolution is used widely. The *stride* is the size of each convolution step — that is, the number of input points that the kernel traverses at each step. Finally, *padding* is a frame of zeros that can be put around the input. At times, a kernel of a given size that moves with a given stride can end up partially outside input borders, causing computational errors. To avoid this, the input can be padded with zeros at no cost for the result, as the product between padding zeros and the corresponding kernel values will be zero.

### 2.3 Core Concept Three: Error-Driven Learning

Given the concepts introduced in previous sections, it is possible to say that every layer in any neural network takes in something and transforms it into something else. The exact shape of this transformation depends on the weights, such that a same input is transformed into different things when multiplied with different weights. For example, consider the vector **x** = [0.27, 0.79, 0.18]. Using no activation functions, multiplying **x** by matrix:

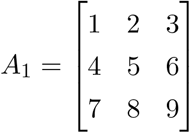

yields the vector **a**_**1**_ = [2.39, 6.11, 9.83]. However, multiplying it by matrix:

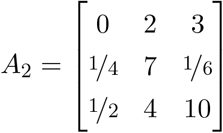

yields the vector **a**_**2**_ = [2.12, 5.62, 5.09] (truncated to the second decimal digit). Therefore, we face two interrelated facts:

1. The output of an ANN is shaped by the weights: when weights change, so does the output (Figure 4)
2. If we want a network to produce a given output, we need to tune the weights

**Figure 4.**
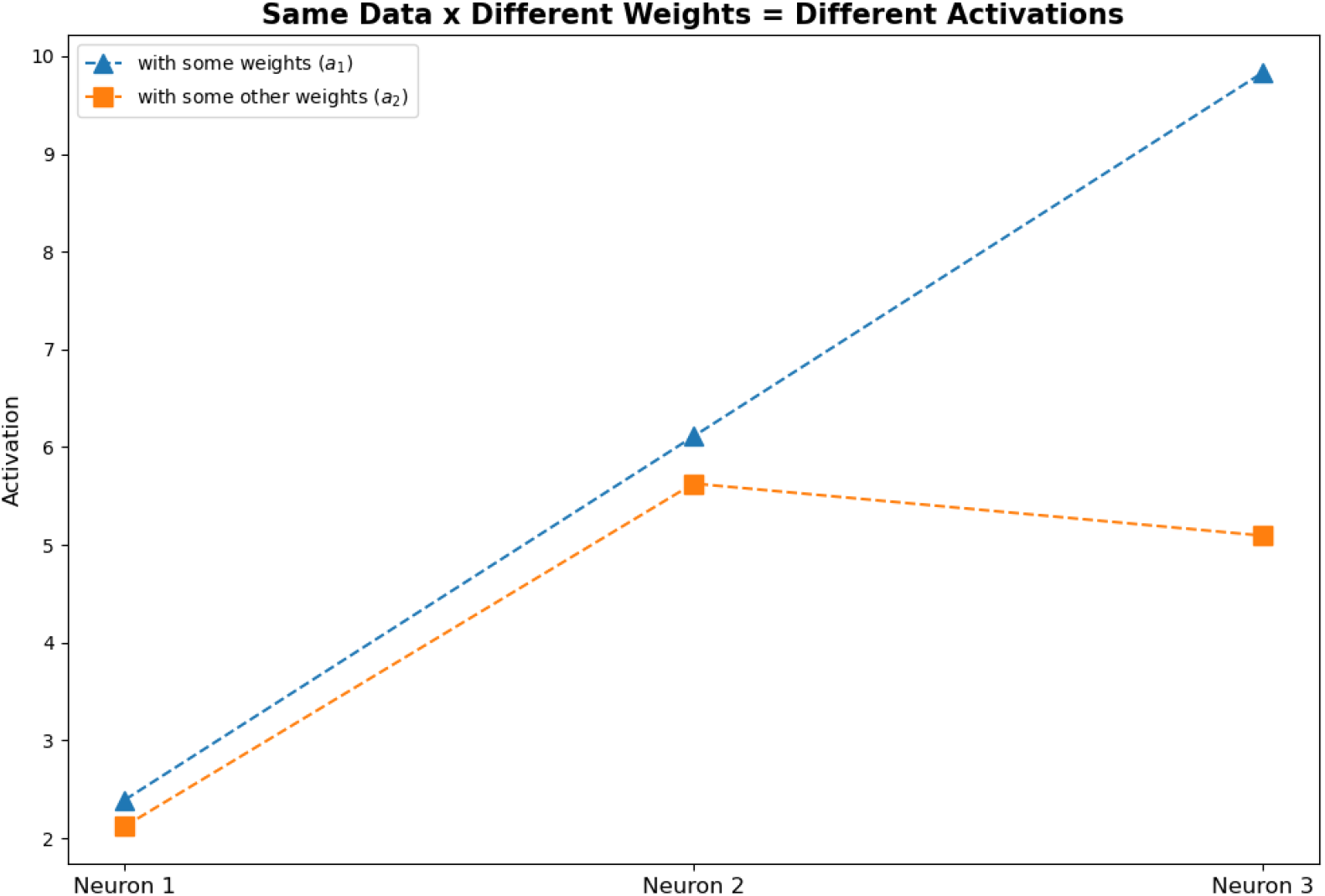
When weights change, activations can change dramatically even if data stay the same

These facts introduce a problem: how can we change the weights towards a desired output? The most common (and simplest) strategy to solve this problem is *error-driven learning*. Error-driven learning is a simple algorithm (that is, a sequence of actions) that includes five steps:

1. Feed data to the ANN, obtaining an output in return
2. Compare the output to a target
3. Calculate an error measure (e.g., the difference between the actual and desired outputs)
4. Change the weights
5. Repeat until the error tends to zero

In order to work, error-driven learning must follow three rules:

1. Weights must change in a way that minimises the error, because that is the desired outcome of the process
2. Different weights must change differently, because some contribute to the error more than others
3. Weights cannot change at random, otherwise learning might take forever

In light of these rules, error-driven learning needs three things:

1. An *error (or loss) function* to compute the distance between real and desired output
2. A <u>derivative</u> of the error function with respect to the weights, which quantifies the contribution of each weight to the error
3. A rule to change weights appropriately

A common rule to change weights appropriately is **error backpropagation** [12], which modifies each weight by subtracting a small fraction of its contribution to the error:

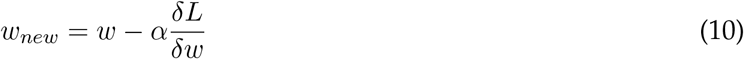

where *w* is any given weight, *α* is the network’s *learning rate* (i.e., an arbitrary coefficient between 0 and 1), and 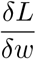 is the <u>partial derivative</u> of loss function *L* with respect to *w* (that is, a measure of *w*’s contribution to *L*). Importantly, the quantity 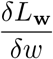 can be positive or negative. If it is positive, it means that *w* is increasing the error and therefore, it should be decreased. Conversely, if it is negative it means that *w* is decreasing the error and therefore, it should be increased.^6^

The rule is called *error backpropagation* because the effect of the error (that is, the weight update) propagates *backwards* through the network, as errors are computed from the network’s outputs and used to update the weights of the last layer, of the one before last, of the second before last… Up until the first. In this context, the output layer shall be thought of as the network’s *front* because it is the endpoint of the information flow through the network (that is, where the flow is headed), while the input layer shall be thought of as the network’s *back* because it is the starting point of the flow. For this reason, the flow of information from data to output is often referred to as *forward pass* (back-to-front), while the flow of information from error to first-layer weight updates is referred to as *backward pass* (front-to-back).

### 2.4 Core Concept Four: Training, Validation, and Testing

An algorithm is valuable if it produces a desired output consistently. Therefore, error-driven learning must modify weights in a direction that allows the network to acquire strong knowledge about the domain of interest — for example, recognising a cat every time an image contains a cat.

Any type of knowledge is strong when it is based on the intrinsic, defining features of the data. Therefore, strong knowledge is robust to small, context-dependent changes, allowing a learner to recognise a cat regardless of it being brown, black or spotted. Strong knowledge is also independent of spurious covariates, allowing a learner to recognise a cat because it is a cat, and not because it is often close to a cup of milk.

To acquire strong knowledge, ANNs should meet three conditions. First, they should be exposed to a *large* body of data, because the more they see, the more they can learn. Second, they should be exposed to a *rich* body of data, because not all cats are brown. Third, they should generalise their knowledge to novel inputs that are similar (but not equal) to previously processed ones.

To meet all three conditions, ANNs usually undergo three phases, called *training, validation*, and *testing*:

1. During training, an ANN does error-driven learning on a body of input data that is hopefully large and rich
2. During validation, the ANN is fed some previously unseen samples, but the weights are not updated (i.e., no learning occurs). This can be done at various stages and with various goals — for example, every *n* training iterations with the goal of obtaining a sanity check for the ongoing training process
3. Finally, the ANN does testing. Here, it is fed previously unseen samples from a separate dataset, without any change of weights. This is the moment of truth, when it is possible to verify if the algorithm has acquired robust knowledge in the domain of interest

It is important to underline that training, validation and testing are carried out on thee different sets of data which come from the same domain (for example, cat pictures), but fulfil different needs (Figure 5). The training set contains data that are only used for *learning* — that is, repeatedly process all samples in the set and update weights after every repetition, until the weights are such that the error is minimum. The validation set contains data that are used to *estimate* test performance before the actual test. Finally, the test set contains data that are used to *deploy* the model — that is, use the knowledge acquired during training to solve the problem at hand, potentially forever.

**Figure 5.**
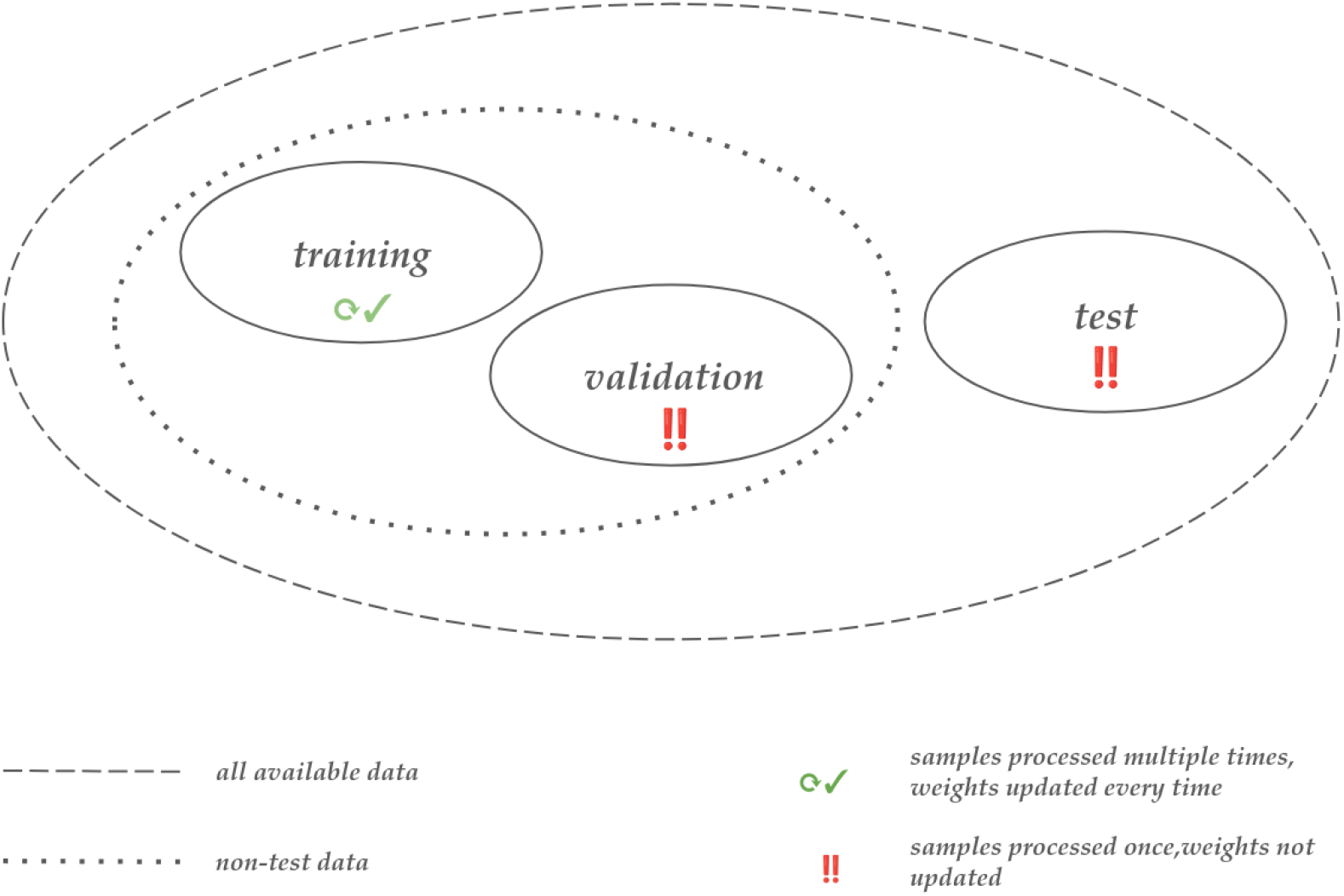
Training, validation, and test sets

Among the three phases, validation is perhaps the most complex to treat and understand. This is due to different people giving validation different flavours, depending of the structure of their dataset and the application scenarios that they face. In particular, one might find the following situations:

- When the training and test sets are large, validation is merely a sanity check for training. In fact, this is a setting where the training set is large enough to yield good learning and the test set is large enough to support strong inferences
- When there is no designated test set due to practical constraints, the validation set serves to estimate what would happen in an actual test. This is the case of ANN models that deploy in real-world settings (for example, in industry or healthcare), where the test set are the samples that will be found in the real world during deployment and are therefore not available during model development
- When the designated test set is small or otherwise biased, the validation set serves as an aid to estimate the true power of the model, as the test set alone does not suffice due to insufficient size
- Finally, the validation set can be used for *model selection*, also known as *hyperparameter tuning*: a complex process that deserves a dedicated section

#### 2.4.1 Model Selection, or *Hyperparameter Tuning*

Learning in a neural network is a complex process that is influenced by a myriad different factors, including the number of layers, their size, the learning rule^7^, or the value of the constants in the weights update rule (e.g., the learning rate). All the elements that are not a network’s weights but can influence learning are called *hyperparameters*. Each possible combination of hyperparameter values defines a different neural network (or *model*) with a different learning potential, so it is vital to identify and use the optimal one. The process of finding the optimal combination of hyperparameter values is known as *model selection* in the statistics literature and *hyperparameter tuning* in modern machine learning practice.

There usually exists an infinite number of possible hyperparameter combinations, and it is not humanly possible to try them all. Therefore, researchers must use some criterion to select a finite number of candidates. Two common strategies are *random search* and *grid search*. In random search, researchers define a search space for each hyperparameter of interest (for example, all the numbers in a given range) and values are sampled at random from that space. In grid search, researchers manually define lists of candidate values for each hyperparameter on the basis of previous knowledge about which values might work. In both cases, the result is a set of combinations that contain one candidate value for each hyperparameter of interest. Each combination is used for a short training-validation cycle, then the combination that yields the lowest validation error (that is, the best-performing model) is selected for the actual training.

The exact structure of the training-validation cycle used for a random or a grid search can vary a lot. One popular strategy is to do a *k-fold cross-validation*, which involves splitting non-test data into *k* subsets (or *folds*), training on *k* − 1 folds, validating on the remaining fold, and repeating the process using a different fold for validation. When used for model selection, k-fold cross-validation translates into an algorithm like the following:

~~~
for each model
    for each permutation of k non-test data subsets
         train on k-1 subsets, validate on the remaining one
          save the validation error
    compute the average validation error
~~~

This is illustrated in Figure 6, where the dataset is partitioned into *k* = 10folds as customary. Other common values for *k* are *k* = 5 and *k* = *N*, where *N* is the size of the dataset [**?**] ^8^.

**Figure 6.**
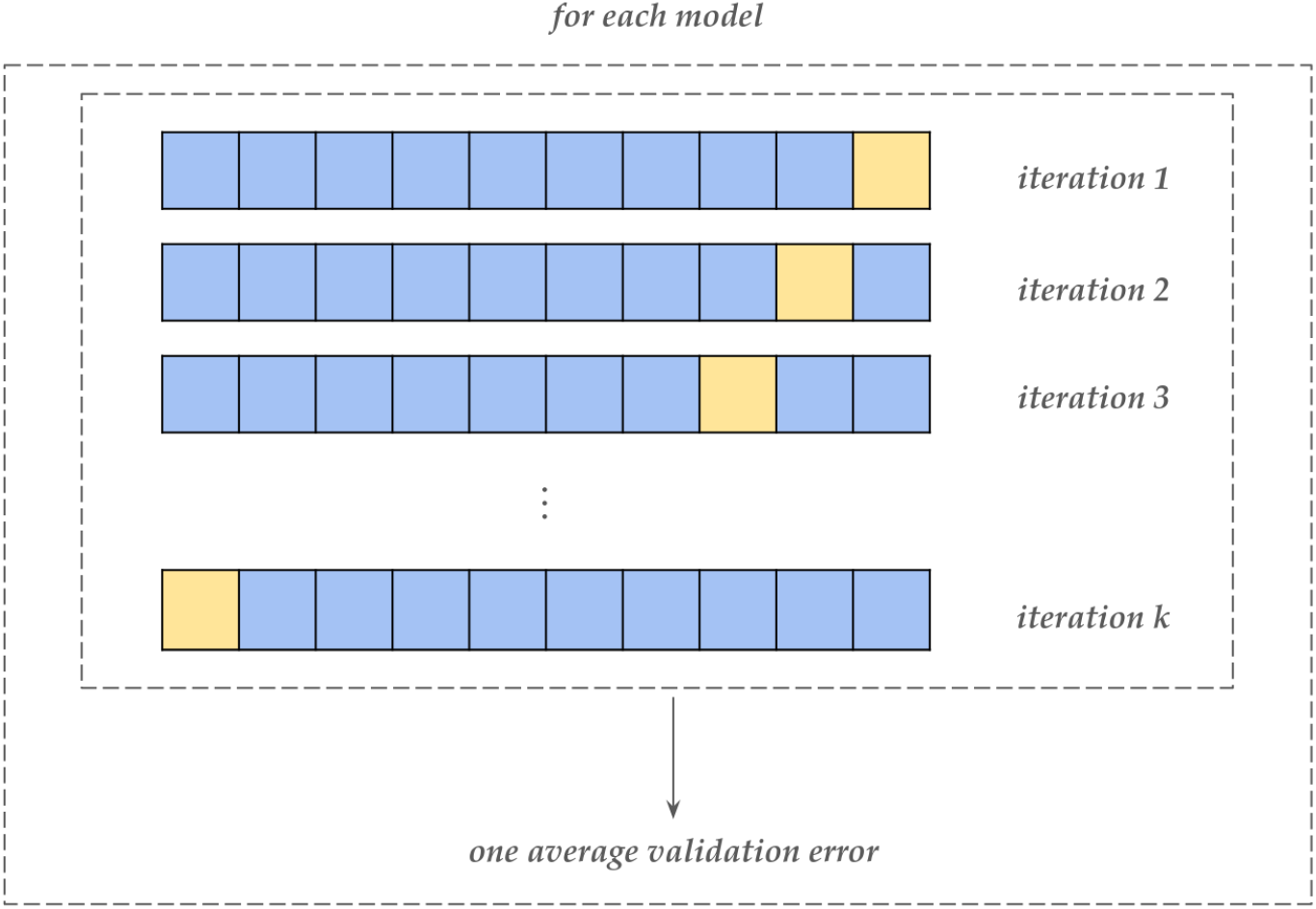
k-fold cross-validation with *k* = 10. The large rectangles represent the dataset, which is partitioned into *k* = 10folds represented by the smaller squares. Blue folds are used for training, while yellow folds are used for validation. At iteration one, the first 9 folds are used for training and the 10^*th*^ is used for validation. At iteration two, the first 8 folds and the 10^*th*^ one are used for training, while the 9^*th*^ is used for validation. At iteration three, the first 7 folds, the 9^*th*^ and the 10^*th*^ are used for training, while the 8^*th*^ is used for validation. This process goes on until the 10^*th*^ (that is, *k*^*th*^) iteration, when the 1^*st*^ fold is used for validation and the remaining 9 for training. At every iteration, the model produces a different validation error, which is saved. At the end of the process, all validation errors are averaged and the average value is taken as a reasonable estimate of the performance that the model would produce at test. The model that produces the least average validation error will thus be selected for training and a subsequent test.

Whatever the value of *k*, the role of cross-validation is to estimate the performance that a model will produce at test in a way that is robust to random sampling of the dataset. In fact, sticking to a single choice of the training and validation sets creates a risk of underestimating model power because either set might contain samples that are particularly easy or hard, biasing the validation error. Conversely, training and validating on multiple different subsets exposes the model to a variety of samples in both the training and validation sets, increasing the robustness of its performance metrics.

While it can increase the robustness of a model and the credibility of one’s work, cross-validation is not compulsory. When the dataset is large the likelihood of incurring in a biased validation set is reasonably low, so it is possible to avoid cross-validation and spare computational resources^9^. Cross-validation (and, more generally, validation) is mostly suggested when the test set is small or unavailable. In such cases, validation or cross-validation are used to increase the number of samples that can be used to estimate test performance [33] . The next section of this tutorial will use a dataset that is relatively easy and relatively large, with 60000 training samples and 10000test samples. In this case — and in many other deep learning scenarios — cross-validation is not a strict requirement.

## 3 The Implementation

Previous sections have introduced the theoretical minimum required to understand ANNs and CNNs, with code examples to complement the explanations. Such examples described the basic machinery behind neural networks, but they were much simpler than actual use cases — which might include high-dimensional inputs, deep networks with different layer types, or complex activation functions. Despite dealing with simplified scenarios, the code examples got more and more complex throughout the paper — and they would be even more complex in a real-world scenario. To navigate such complexity without drowning into implementation details, researchers need software that lets them focus on high-level design choices while maintaining full control of all scientific aspects. In 2026, such software is available in all the three major scientific programming languages: Python, R, and Matlab. While it is hard to prove it quantitatively, Python seems to offer the highest-quality, most used, and most maintained solutions through the PyTorch [17] and TensorFlow [34] frameworks — which are free, open source, and widely used across scientific and industrial settings. The two frameworks have different strengths and weaknesses, but they are largely equivalent from the perspective of an entry-level user. However, PyTorch has a tight integration with NumPy and is the cornerstone of several softwares of academic interest, like [PyTorch Geometric](https://pytorch-geometric.readthedocs.io/en/latest/) [35] for graph neural networks or [Braindecode](https://braindecode.org/stable/index.html) for decoding brain electrophysiological data [26]. Therefore, this paper and the [associated Jupyter Notebook](https://github.com/matteo-d-m/cnns-tutorial) propose PyTorch-based implementations.

### 3.1 Enter: PyTorch

PyTorch is a Python framework to access publicly available machine learning datasets, program ANNs, and exploit the parallel computing capabilities of NVIDIA graphics processing units (GPUs), which can greatly accelerate computations compared to the usual central processing units (CPUs). This last point is one of PyTorch’s main strengths, as computational resources can be a major pitfall in developing ANNs. PyTorch can exploit the computing power of GPUs thanks to its tight integration with CUDA, a software package to perform general-purpose computing on GPUs (which would otherwise be dedicated entirely to graphics). The usefulness of CUDA is increased by cuDNN, a library of hardware-efficient implementations of operations that arise frequently in deep neural networks, such as convolution and pooling. While very powerful, CUDA and cuDNN can be difficult to use for non-field-experts. Therefore, PyTorch provides a Python interface that allows everyone to use CUDA and cuDNN in the absence of domain expertise. Importantly, PyTorch is free and open source — like the Python programming language and most Python-based tools.

Users that do not want to perform any installations can use PyTorch in [Google Colab](https://colab.research.google.com/) — a free service where users can edit code in a browser and run it on a remote server without needing any set-up. The [companion Jupyter Notebook](https://github.com/matteo-d-m/cnns-tutorial) for this paper includes an `Open in Colab` button that sends straight to the service. Users that feel confident installing things on their computer can follow the instructions on [PyTorch’s official website](https://pytorch.org/). As of January 2026, the process boils down to opening a terminal and running Code Example 6:

#### Code Example 6: Installing PyTorch on Linux or Windows (January 2026 — CUDA version 12.6)

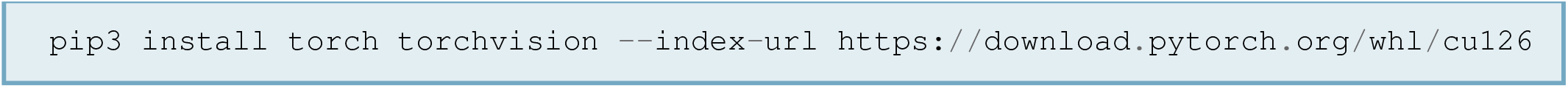

As in any other Python workflow, it is highly advisable to install PyTorch in a dedicated `conda` environment. `conda` environments are *“self-contained, isolated spaces where you can install specific versions of software packages, including dependencies, libraries, and Python versions. This isolation helps avoid conflicts between package versions and ensures that your projects have the exact libraries and tools they need”* ([source](https://www.anaconda.com/docs/tools/working-with-conda/environments) – accessed on 27 January 2026). To this end, the installation of PyTorch must be preceded by an installation of the Anaconda package manager, preferably through the lightweight Miniconda distribution. All instructions and relevant information can be found on the Anaconda documentation website at [this](https://docs.conda.io/projects/conda/en/stable/index.html) link.

### 3.2 A Case Study

The following sections will demonstrate every PyTorch core component, using a simple image recognition task as case study. All the demo code is contained in the [companion Jupyter Notebook](https://github.com/matteo-d-m/cnns-tutorial), which can be run on Google Colab as well as in a local set-up.

#### 3.2.1 Datasets & DataLoaders

To perform image recognition, one needs images. PyTorch includes a library called `torchvision` which, among other things, contains a `datasets` package that provides tools to automatically download and use publicly available image datasets. One classic example is the MNIST database of handwritten digits [36], which contains 70000handwriting samples of the numbers from 0to 9, encoded as 28 x 28 grids of gray-scale luminosity values (Figure 7).

**Figure 7.**
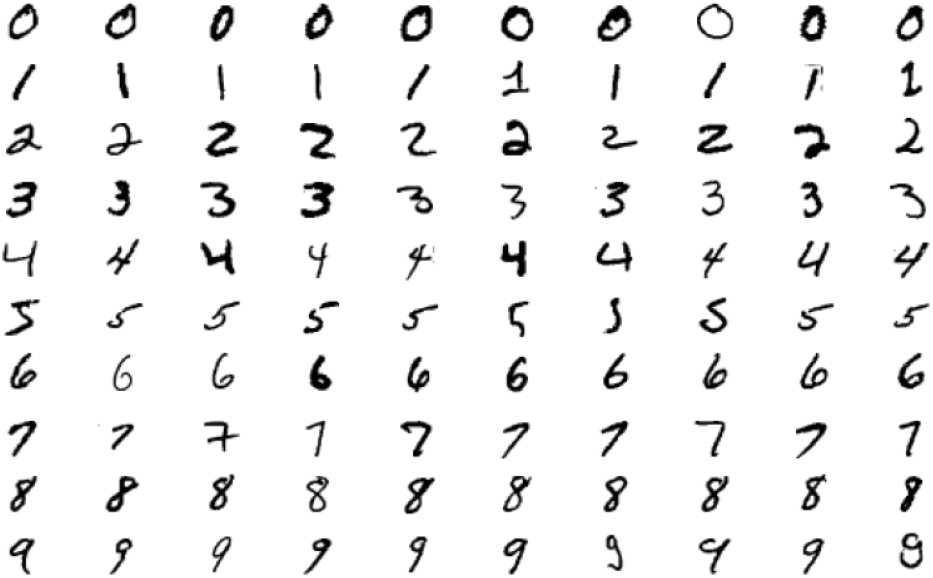
Samples from the MNIST database [36]

Like many other datasets, MNIST can be downloaded directly in PyTorch with a single command. The resulting variables are `Dataset` objects, which represent a set of data and include two crucially important methods (that is, operations). The first one is `__len()__`, which allows a user to get the dataset’s length, and the second one is `__getitem()__`, which allows a user to index into the dataset and access samples at will. The double underscores before and after the name of those method mean that they are *magic* methods — that is, users do not call them explicitly, but the methods work under the hood when users call `len(dataset_name)` and `dataset_name[index]`, respectively.

Code Example 7 downloads MNIST from its online repository, doing two separate downloads for the training and test sets. Subsequently, Code Example 8 uses `__len()__` to print the number of samples contained in the training and test datasets. The corresponding output would be that the training dataset contains 60000 samples, while the test dataset contains 10000. Finally, Code Example 9 uses `__getitem()__` to extract an arbitrary sample (for example, sample number 0) and familiarise with it.

##### Code Example 7: Downloading MNIST with PyTorch

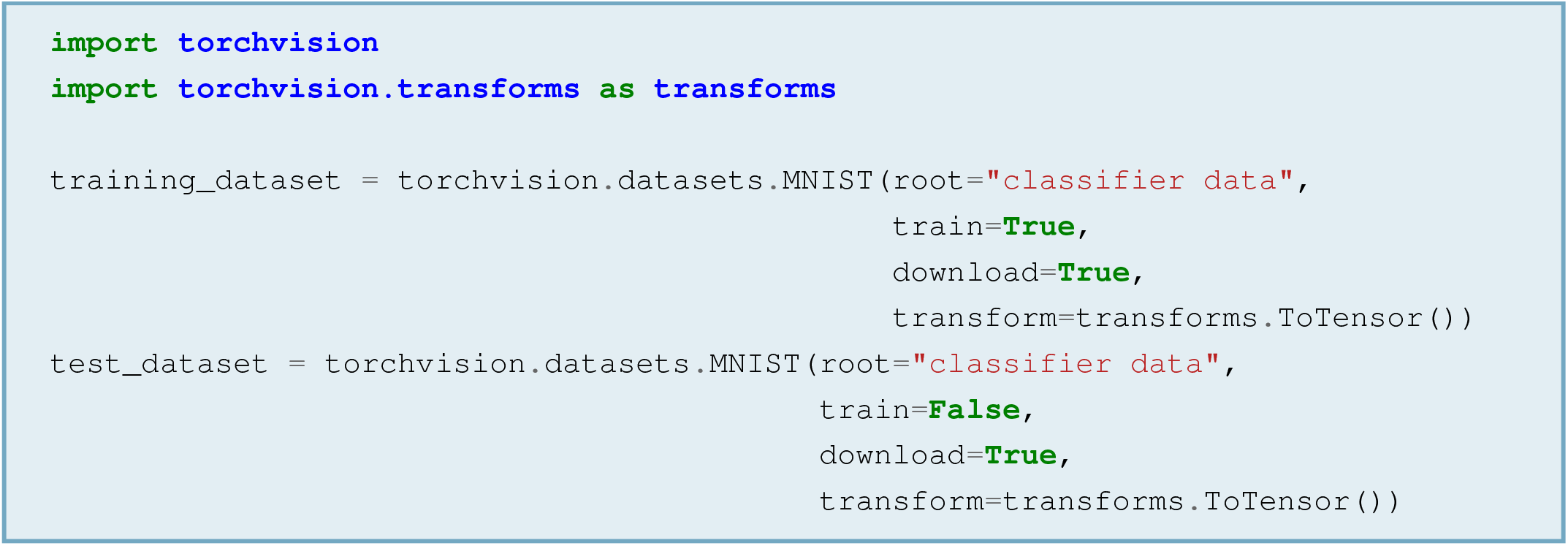

##### Code Example 8: Getting the Length of a Dataset Object

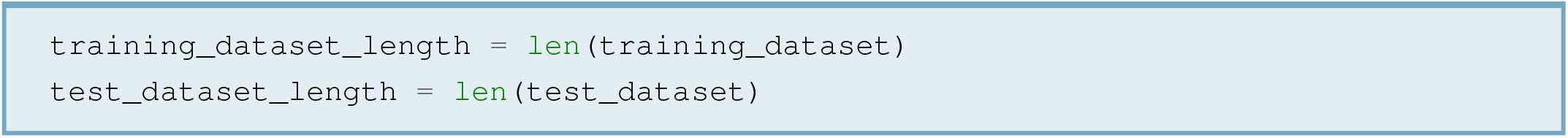

##### Code Example 9: Extracting a Sample from a Dataset Object

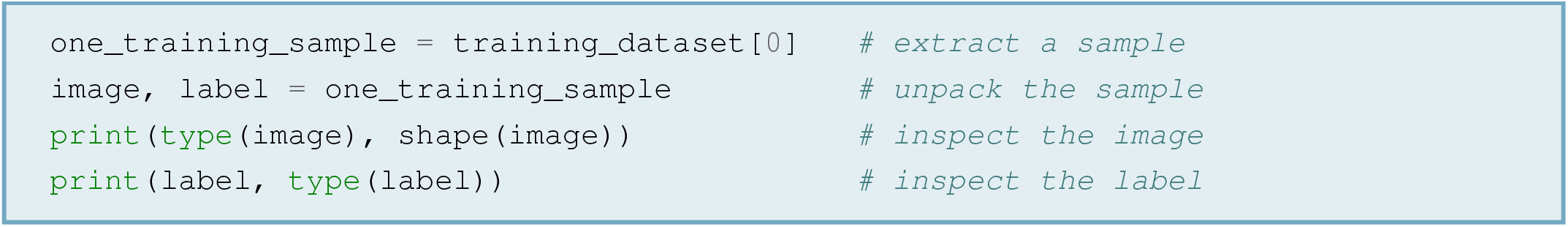

The output of Code Example 9 would be that each training sample (or test sample, for that matter) is a tuple (that is, an ordered sequence) of two elements: an image and a label. The image is encoded as a tensor, which is the central data structure of PyTorch and is basically an array with an arbitrary number of dimensions. Neuroscientists that have worked with neuroimaging data have seen tensors under a wealth of other names: for example, epoched EEG data are three-dimensional tensors (`epochs x channels x time`), and magnetic resonance images are four-dimensional tensors (`slices x width x height x depth`). PyTorch tensors are the same as standard arrays, except that they can be easily loaded into GPUs and they are optimised for fast, automatic differentiation of the functions that they represent — which is vital for backpropagation^10^. Like standard NumPy arrays, PyTorch tensors have a `.shape` attribute, which provides information about the tensor’s shape. In the case of MNIST images, tensors have shape `1 x 28 x 28` because they store images with 1 colour channel and 28 x 28 shape. The fact that MNIST images have *one* colour channel means that they are black and white. Were they coloured, they would be a combination of a red (R), a green (G), and a blue (B) image, so they would have 3 channels: one that stores the R part of the image, one that stores the G part of the image, and one that stores the B part of the image.

Once familiarised with the dataset, one would typically reserve some data for validation. As explained before, there is a wealth of strategies for doing that (see Core Concept Four). The simplest one is to reserve a randomly chosen fraction of the training data (for example, 25% of the total) for validation and use the rest for training. To this end, PyTorch provides a function called `random_split()` (from the `data` module of the `utils` package), which slices a dataset into *n* subsets of desired length. Code Example 10 uses it to reserve 25% of training data for validation, leaving the remainder for the actual training.

##### Code Example 10: Training-Validation Split

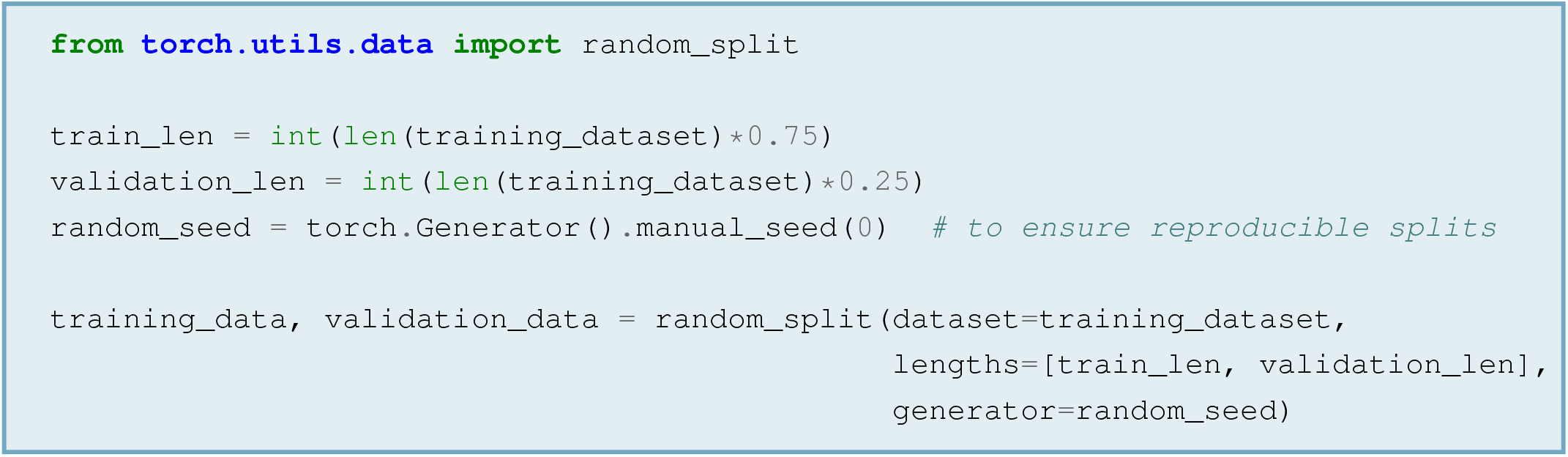

Whether used for training, validation or testing, `Dataset` objects can be iterated over — that is, it is possible to process all their elements sequentially. Unfortunately, `Dataset` objects behave like standard Python iterables, which means that samples can only be accessed one at a time. However, when training ANNs it is desirable to access multiple samples at once and process them as if they were a single sample: a process known as *batch* (or *mini-batch*) *processing*, which can decrease computation time dramatically. It is also desirable to exploit all the available computational power and to shuffle training samples at every iteration. To achieve these goals, PyTorch provides users with `DataLoader` objects. A `DataLoader` object takes a `Dataset` object and empowers it with batch processing capabilities (that is, organise samples in batches that can be processed as one), multiprocessing capabilities (that is, use all the computational power provided by the machine) and shuffling capabilities (that is, access samples in a different order at every training iteration, to avoid possible order effects on learning). Code Example 11 creates one `DataLoader` for each of the training, validation, and test datasets. Shuffling is enabled for training only, because order should not be an issue during validation and testing.

##### Code Example 11: Initialising DataLoaders

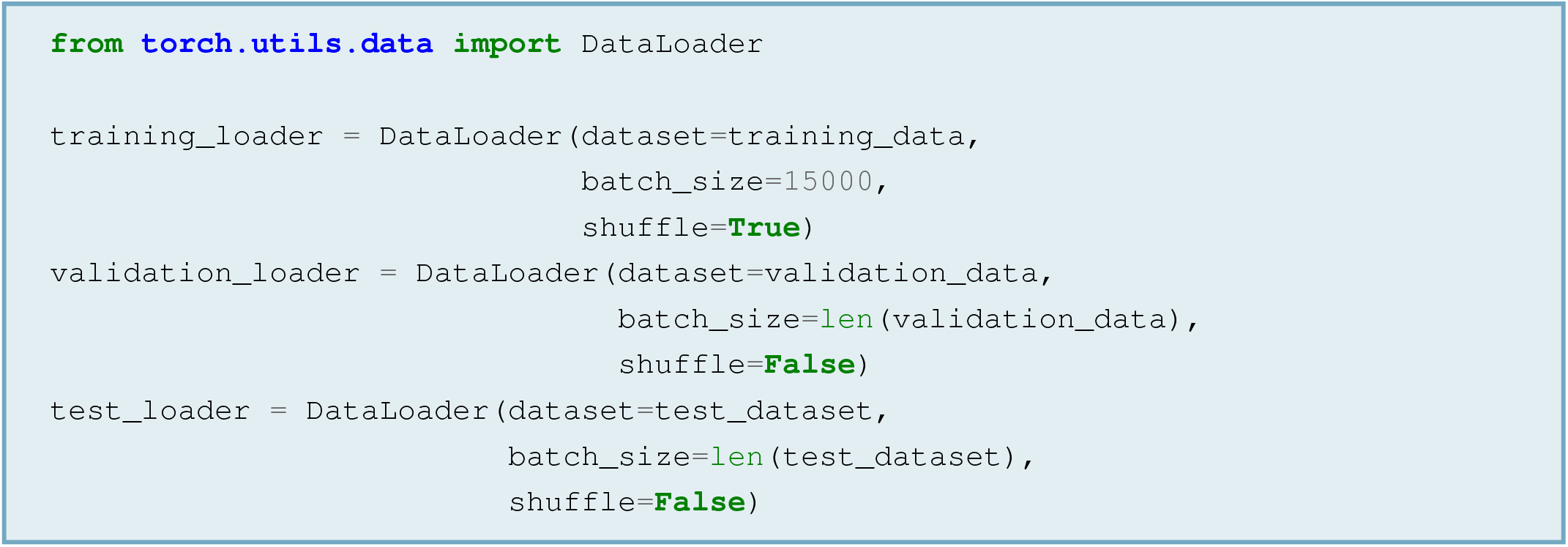

Like `Dataset` objects, `DataLoader` objects implement a `__len()__` method. Therefore, it is possible to print and use their length as done with datasets. However, dataloaders have no `__getitem()__` method, so running `dataloader[index]` will return an error message saying that dataloaders are “not subscriptable” — that is, they cannot be inspected on a sample-by-sample basis. Nonetheless, `DataLoader` objects can (and should) be iterated over. By using this opportunity, it is possible to get a deeper sense of their contents by printing the properties of samples (for example, their type and shape):

##### Code Example 12: Inspecting a DataLoader

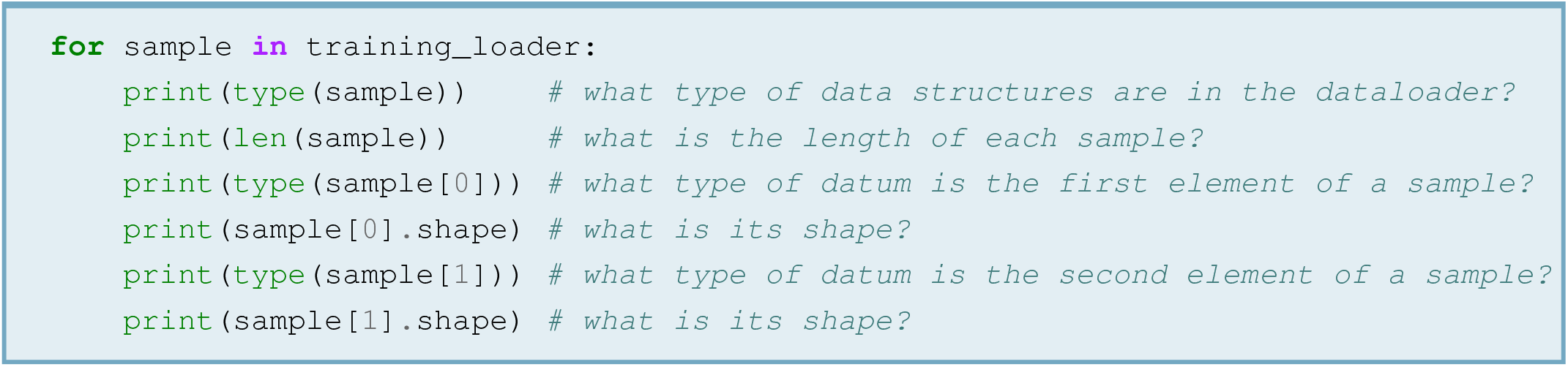

The outputs of the code above would reveal that the training loader contains three items (one per batch), each of which is a list of two elements: a tensor of shape `batch size x channels x height x width` (that is, a batch of images) and a tensor of shape `batch size` (that is, a batch of labels). The items in the dataloader are three and have length 15000 because the training dataset contains 45000 items (that is, 75% of the 60000 found before the training-validation split) and the batch size was set to 15000, which is a third of 45000. During training (and, similarly, during validation and testing), batches will be fed to neural networks as if they were a single tensor, using a syntax of the type:

##### Code Example 13: Batch Processing Pseudocode

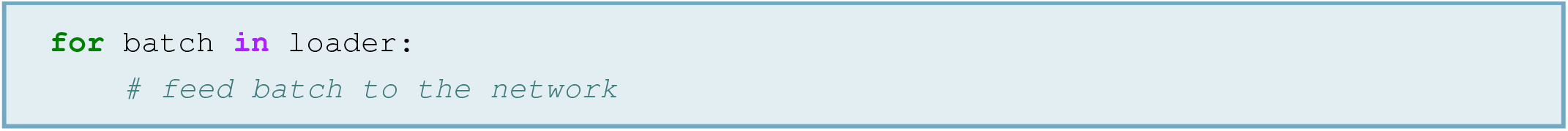

#### 3.2.2 Building a CNN

Having obtained a dataset and an efficient way to access samples, it is time to build a CNN that could learn them. In PyTorch, neural networks are implemented as **classes** — that is, Python entities defined by attributes and operations to manipulate such attributes. This makes sense in light of two simple facts:

1. A neural network has structural components, which are its layers. These are the *attributes* of the neural network class — in other words, its building blocks
2. A neural network performs a chain of operations on data. These are the *methods* of the neural network class — in other words, its capabilities

Code Example 14 implements a deep CNN with two convolutional layers followed by standard, non-convolutional layers:^11^

##### Code Example 14: A Simple CNN

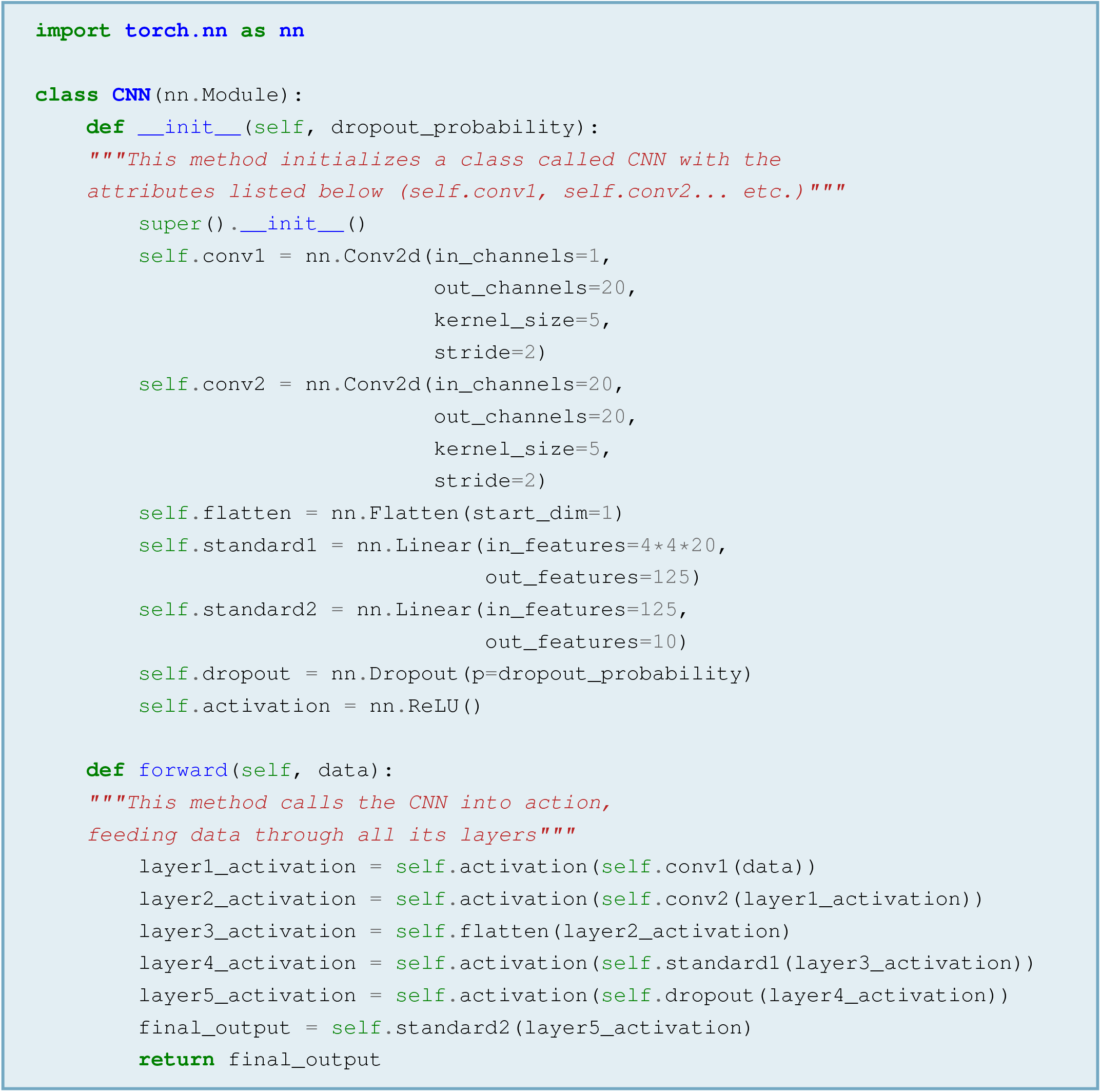

The code above includes some new terms from the PyTorch jargon:

- **Input channels** (`in_channels`). As mentioned before, image and other types of data can be elementary (like black-and-white images) or they can be a combination of multiple sub-inputs (like RGB coloured images). When the input is elementary, the convolution is said to have one input channel. When the input is composite, the convolution is said to have multiple input channels. In case of multiple input channels, the convolution kernel operates over each channel and the activations resulting from each convolution step are summed to obtain a single feature map for output
- **Output channels** (`out_channels`). This is the number of feature maps that the convolution layer should produce. Until now, this paper has considered the simple case of a single kernel that produces a single feature map. However, real-world CNNs tend to use multiple kernels that result in multiple feature maps. There are at least two reasons why it is desirable to use multiple kernels, of which one is neuroscientific and the other technical:
  – The neuroscientific reason is that multiple kernels generate multiple feature maps, simulating distributed processing in the brain. In fact, each map can be seen as representing an activity pattern over a set of neurons — akin to a single neuronal ensemble reacting to a sensory input — while multiple maps can be seen as representing multiple activity patterns over different sets of neurons, akin to multiple neuronal ensembles reacting to the same input at once
  – The technical reason is that before the start of training, convolution kernels are initialised according to some heuristic that sets initial values for the weight parameters [37]. Such values have little in common with the values they should have after learning, but set a starting point to their evolutionary path — that is, the trajectory of change that weights will undergo during learning. Some starting points may be better than others — that is, they may require a smaller number of weight updates for the network to achieve an optimal performance. Therefore, using multiple kernels increases the probability of setting an optimal start to the learning process
- **Flatten** (`nn.Flatten`). After the last convolution, all feature maps must be collapsed into a single vector for use with standard layers. To this end, each feature map is reshaped into a one-dimensional vector (in other words, it is *flattened*) and the results are concatenated into one big vector. This is done by the `Flatten` class contained in the `nn` package
- **Linear** (`nn.Linear`). This term indicates a standard neural network layer, emphasising that it operates a linear transformation of the input. Other synonyms — like *fully connected* and *dense* — emphasise that every element of the input contributes to every element of the output (cf. 2.1.4)
- **Input features & Output features** (`in_features`, `out_features`). Input features are the number of elements contained in the input to a linear layer, while output features are the number of elements that should be contained in the output. Note that:
- The input features are dictated by the size of the input, be it external data or the output of previous layers. In the code example, the second convolutional layer outputs 20feature maps, each one 4 x 4. This means that after flattening, there will be a vector containing 20· 4 · 4 = 320elements. Therefore, the input features to the first fully-connected layer must be 20· 4 · 4 = 320
- The output features are arbitrary
- **Dropout** (`nn.Dropout`). This is a special type of neural network layer that turns randomly selected weights to zero with a given probability [38]: for example, a dropout probability of 0.25 turn a random 25% of weights to zero. While relatively obscure from the theoretical standpoint, dropout is known to improve learning in a variety of neural network architectures

#### 3.2.3 Model Selection / Hyperparameter Tuning

There are a number of techniques to do hyperparameter tuning in PyTorch, many of which require dedicated external libraries like [Optuna](https://optuna.org/) [39]. However, those are mostly useful for advanced tuning procedures and users that are already familiar with this world. From an educational perspective, it might be better to code a simple search from scratch — for example, a *grid search*.

A grid search is a simple hyperparameter optimization procedure where users define a set of hyperparameters, assign them a list of candidate values, and try a model for all the resulting combinations. For example, one might be interested in hyperparameters *A* — with possible values *a*_1_ and *a*_2_ — and *B* — with possible values *b*_1_ and *b*_2_. This would result in four possible combinations: *{*(a_1_, *b*_1_), (a_1_, *b*_2_), (a_2_, *b*_1_), (a_2_, *b*_2_)}, corresponding to four different model specimens. Each model specimen would be tried for a number of iterations before deciding if its performance is satisfactory enough to select that model for the full trainingvalidation-testing cycle.

Code Example 15 defines a list of candidate values for three different hyperparameters: learning rate, weight decay, and dropout probability. The *learning rate* is the constant that controls the magnitude of weight updates during backpropagation (cf. Equation 10 in section 2.3). *Weight decay* is another constant that controls learning by keeping weights generally small in order to avoid giving excessive relevance to specific features. Finally, the *dropout probability* is the probability that any given weight is set to zero in dropout layers: for example, setting it to 0.25 implies that 25% of the neurons in the affected layer will be forcefully inactivated at any given iteration.

##### Code Example 15: Grid Search – Step 1: Setting Candidate Hyperparameter Values

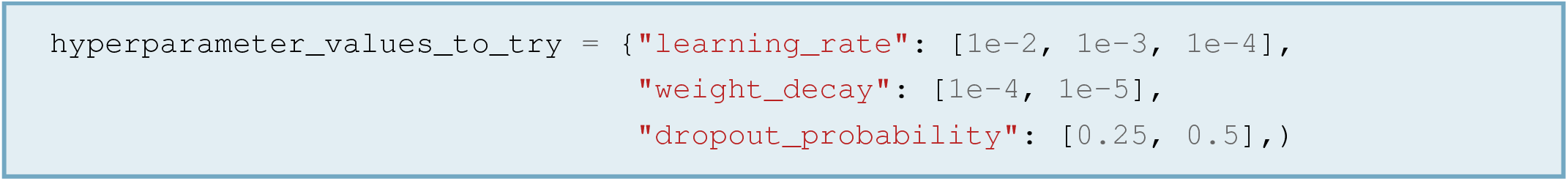

To create a proper grid, the candidate hyperparameter values must be combined through a Cartesian product. The Cartesian product is an operation that, given two or more sets, returns all possible combinations of the items in those sets. Formally, the Cartesian product should return all *ordered* combinations of the available items — that is, it should return (a, *b*) as well as (b, *a*). However, the order of the items is irrelevant in hyperparameter tuning. Therefore, the product should return only one among (a, *b*) and (b, *a*). This is exactly what is done by the `product()` function, which is contained in Python’s built-in `itertools` module. Code Example 16 leverages `product()` to combine hyperparameter values and collect the resulting combinations in a list.

##### Code Example 16: Grid Search – Step 2: Building a Grid

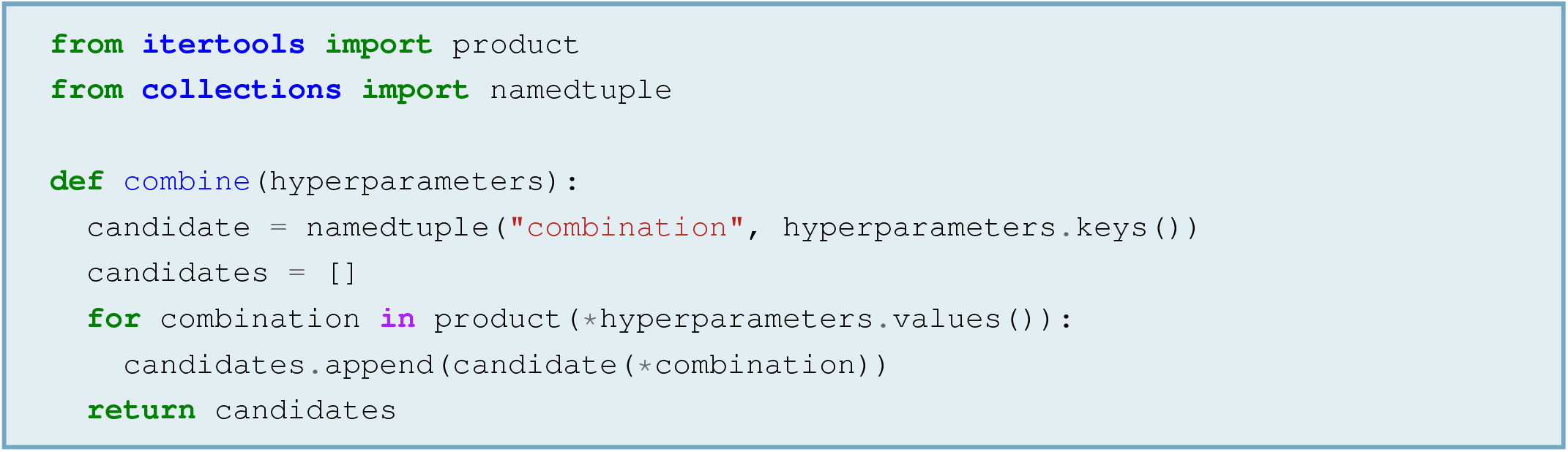

Applying this function to the dictionary defined in Code Example 15 would yield 12 different combinations of hyperparameter values, given by 2 dropout probabilities, 3 learning rates and 2 weight decays.

After building a list of candidate combinations, everything is in place to try them. To this end, one can use a function like the one defined in Code Example 17:

##### Code Example 17: Grid Search – Step 3: The Best-Scoring Model Wins

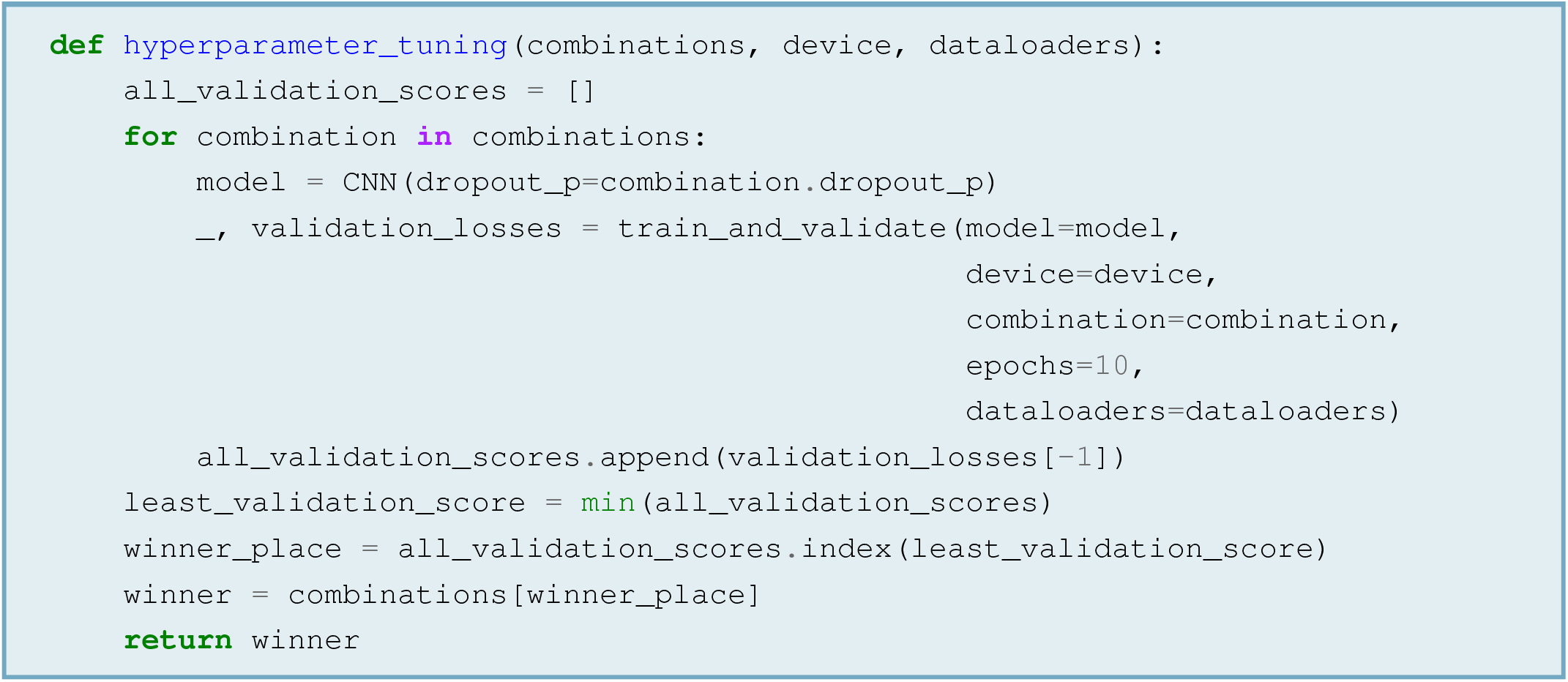

For every candidate combination, this function initialises a CNN with the combination’s dropout probability. Note that this could be any other structural hyperparameter, like the size of a layer or the activation function. After initialisation, the CNN is trained and validated for ten times (`train_and_validate()`). This means that it does a complete pass over the training set for ten times, and at the end of every pass it does another one on the validation set. The result is a list of ten training losses and ten validation losses. The training losses are ignored (_), while the validation losses are saved (`validation_losses`). However, only the last validation loss is kept (`[-1]`), because that one represents the best performance so far. For every model, the validation loss at the 10^*th*^ epoch is saved in a list and subsequently, the smallest value is taken from that list. The combination of hyperparameters that yielded such value (i.e., the one whose position in the list is the same as the position of the minimum in the list of 10^*th*^-epoch losses) is the winning combination, which will be selected for a full training-validation cycle and a subsequent test.

#### 3.2.4 The Training-Validation Loop

We now have a dataset, a CNN, and a hyperparameter tuning method that has returned the most promising CNN specimen. Therefore, we can proceed to a full training-validation cycle using the CNN selected by hyperparameter tuning. The function in Code Example 18 — called `train_and_validate()` — does just that, putting together the pieces introduced thus far. It is the exact same function that was called in Code Example 17 to perform a short training-validation cycle during hyperparameter tuning.

##### Code Example 18: The Training-Validation Loop

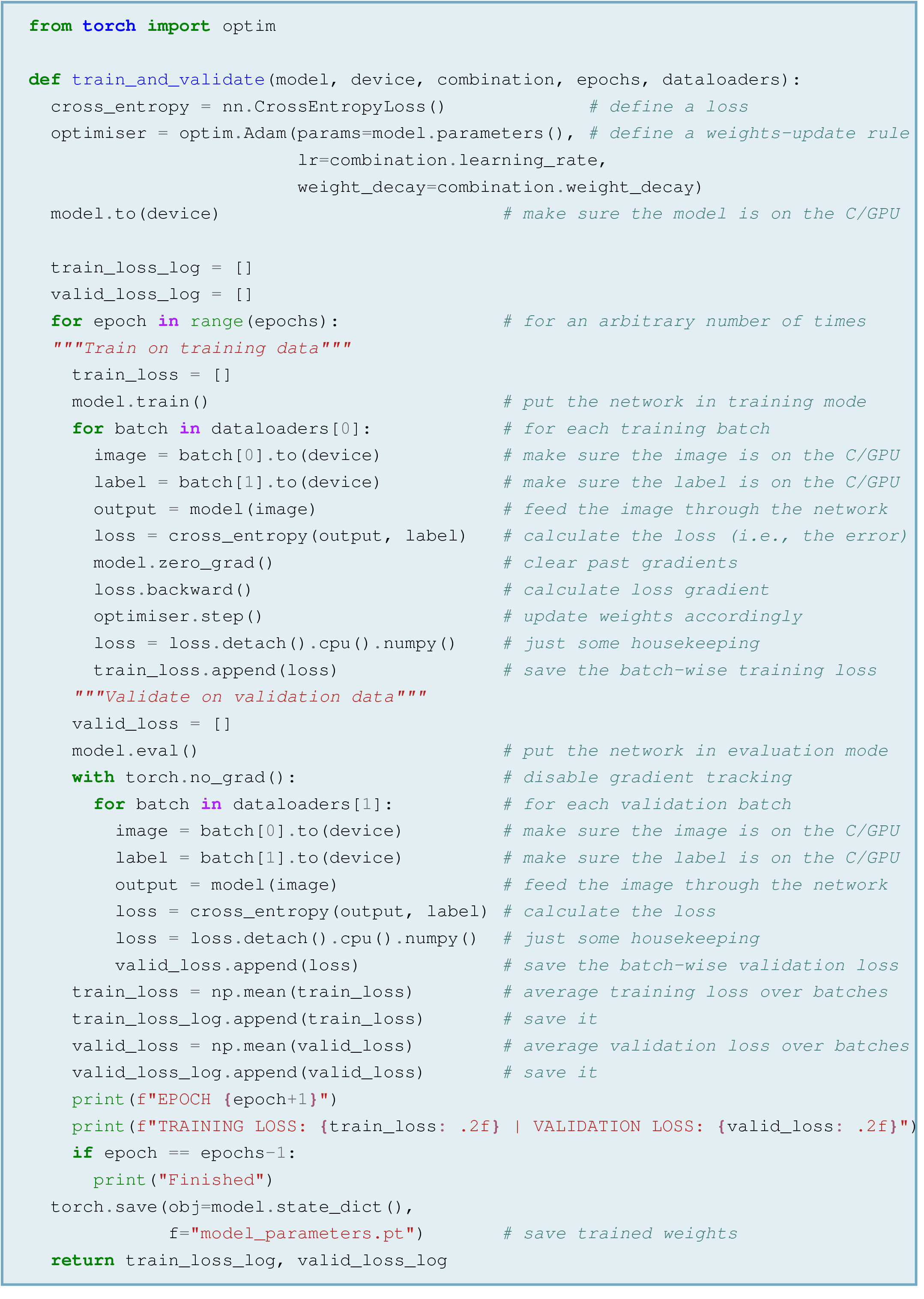

Despite its length, `train_and_validate()` does only a handful of things. First, it defines a loss function using the `CrossEntropyLoss()` class from PyTorch’s `nn` package. <u>Cross-entropy</u> is a measure of the distance between two probability distributions, of which one is a set of labels guessed by the neural network and the other is a set of true labels. As a first step, `nn.CrossEntropyLoss()` rescales the activations from the last network layer so they all belong to the [0, 1] range and they sum to 1 (like probabilities)^12^. The result is a probability distribution over input labels — that is, a set of guesses concerning the identity of the processed data. This set of guesses is compared to the vector of true labels — which contains a 1 for the correct answer and a 0for all others — by means of cross-entropy: the larger the cross-entropy, the larger the distance between the probability distributions, the larger the loss.

After defining a loss function, the code defines an optimiser, which is basically the weights-update rule that will be used for learning^13^ (`optimiser = optim.Adam()`). The optimiser of choice is <u>Adam</u>, a more efficient version of basic backpropagation [40]. To update the weights, Adam needs to know where to find them (`params=model.parameters()`) and what value to use for the learning rate and weight decay, which are both taken from the winning hyperparameter combination (`lr=combination.learning_rate` and `weight_decay=combination.weight_decay`).

Subsequently, the code allocates the CNN to the device that will execute its computations — that is, GPU if available and CPU otherwise.

Having defined everything, the code proceeds to run the actual training. For an arbitrary number of epochs (that is, iterations), the network processes each batch of data. This means that, for each batch, it takes the images and produces tentative labels for output. These tentative labels are compared to the ground truth by means of cross-entropy. Subsequently, the code computes the <u>gradient</u> of the error with respect to the weights, meaning that it estimates the contribution of each weight to the error (`loss.backward()`). This gradient is used to update all weights with the rule specified by the optimiser (`optimiser.step()`), whose conceptual backbone is Equation 10 from section 2.3 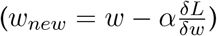. Finally, the batch-wise loss is saved in a list and this ends the training part of the epoch. Next, the network is put to evaluation mode, meaning that training-specific settings (like Dropout) are disabled (`model.eval()`). Without PyTorch keeping track of gradients (`with torch.no_grad()`), the network processes each validation batch. This works exactly as in training, with the only exception that no error gradient is computed and no optimisation step is taken. However, the batch-wise losses are saved in a dedicated list as it happened at training. Subsequently, the code computes the mean loss over batches for both training and validation, saving it in dedicated lists that will end up storing one value per epoch. At this point the training-validation cycle is over: after saving the final value of all weights, the code ends.

**Considerations On The Loss**. Figure 8 shows the loss trend, demonstrating that both the training and validation losses decayed exponentially over epochs. Beside the general trend, there are two interesting facts to note:

1. The first interesting fact is that the validation loss is systematically lower than the training loss for the majority of epochs. One likely reason is that training is subject to *regularisation*, while validation is not. This means that training is intentionally underpowered (in other words, *regularised*) to prevent the A/CNN from *overfitting* — that is, finding weights that are too specific to training data and would not serve well at the validation or test stages^14^. As a result, the performance at training can be worse than the network’s true potential, which shows only at validation. One less likely reason is that the validation dataset might contain easier samples than the training dataset: as the two sets are created with a random split of one father dataset, the easiest samples might in principle end up in one dataset and not in the other
2. The second interesting fact is that the validation loss stops decreasing around epoch 35, while the training loss keeps going down. This means that the network entered the overfitting regime, and any more training would bring more negative effects than benefits

**Figure 8.**
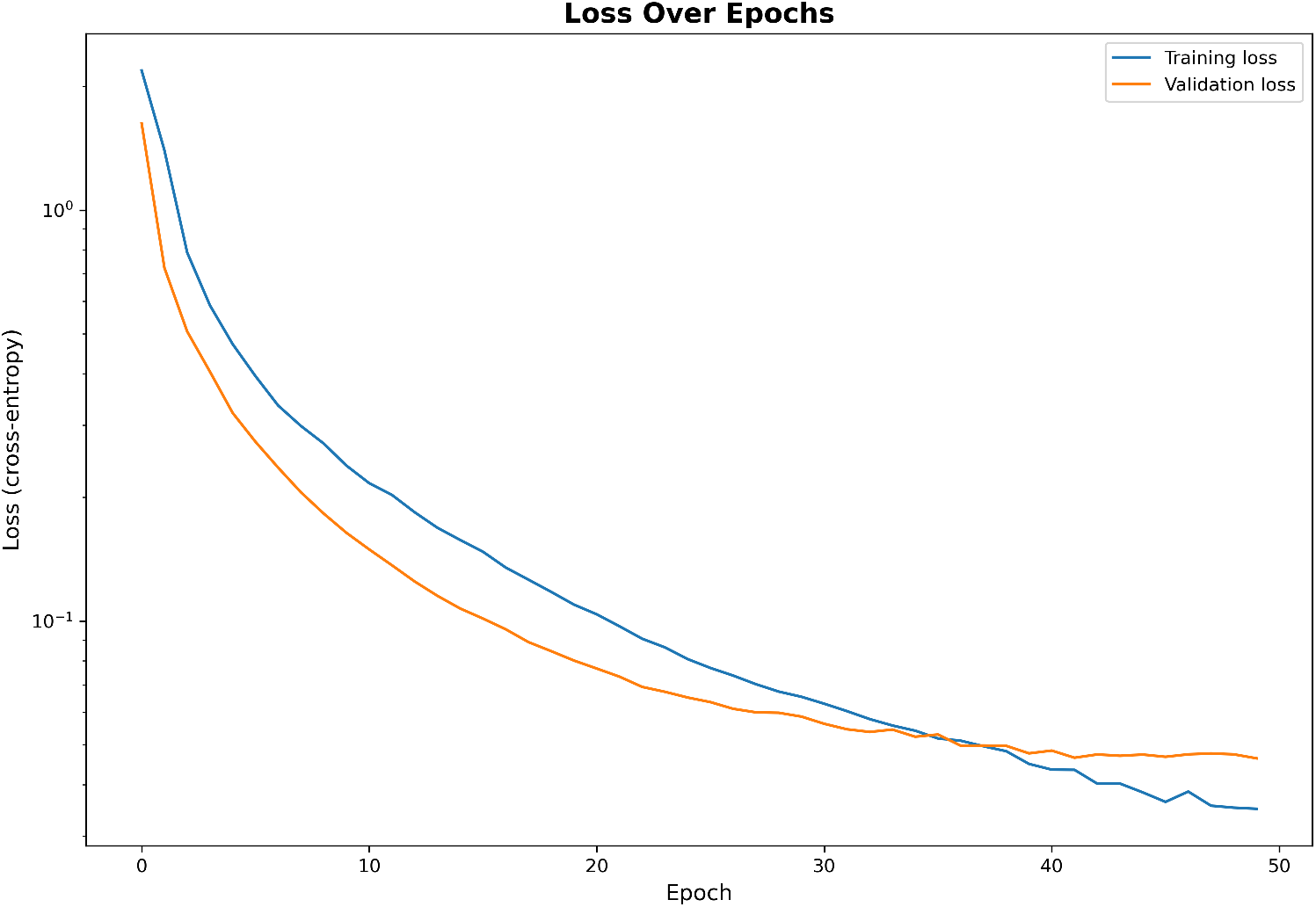
Exponential decay over epochs of the training and validation loss. There are signs of (perhaps excessive) regularisation until around epoch 35 (the orange line being below the blue line), followed by signs of overfitting (the orange line crossing the blue line)

#### 3.2.5 Finally: The Test

Compared to previous steps, the test is rather trivial. As shown by Code Example 19, testing operations are the same as in training and validation — with the only difference that no loss is calculated. Instead, performance is measured in terms of accuracy, which is the percentage of correctly classified samples. For the network presented in this tutorial, the expected accuracy is around 98*/*99%, meaning that the network is expected to output the correct class label for 98*/*99% of test samples.

#### 3.2.6 Inspecting a Trained Model: Basic Techniques

Once trained and tested, a model can be inspected with a confusion matrix. This is simply a matrix that stores the number of correct vs. incorrect responses for each class in the dataset, providing graphical information about which classes were easier vs. harder to learn. The [companion Jupyter Notebook](https://github.com/matteo-d-m/cnns-tutorial) contains code to plot a confusion matrix that looks like Figure 9, where the horizontal axis (i.e., the columns) stores the labels predicted by the network and the vertical axis (i.e., the rows) stores the ground-truth labels. Each entry of the matrix denoted by (i, *j*) stores the number of samples that the network predicted to belong to class *j* when they actually belonged to class *i*: for example, the number in the second row and third column — that is, entry (2, 3) — is the number of times that the network predicted class 3 when the true label was class 2. The implication is that the numbers on the main diagonal (that is, the line from top-left to bottom-right) count the number of correctly classified samples, as the main diagonal is the place where *i* = *j*.

**Figure 9.**
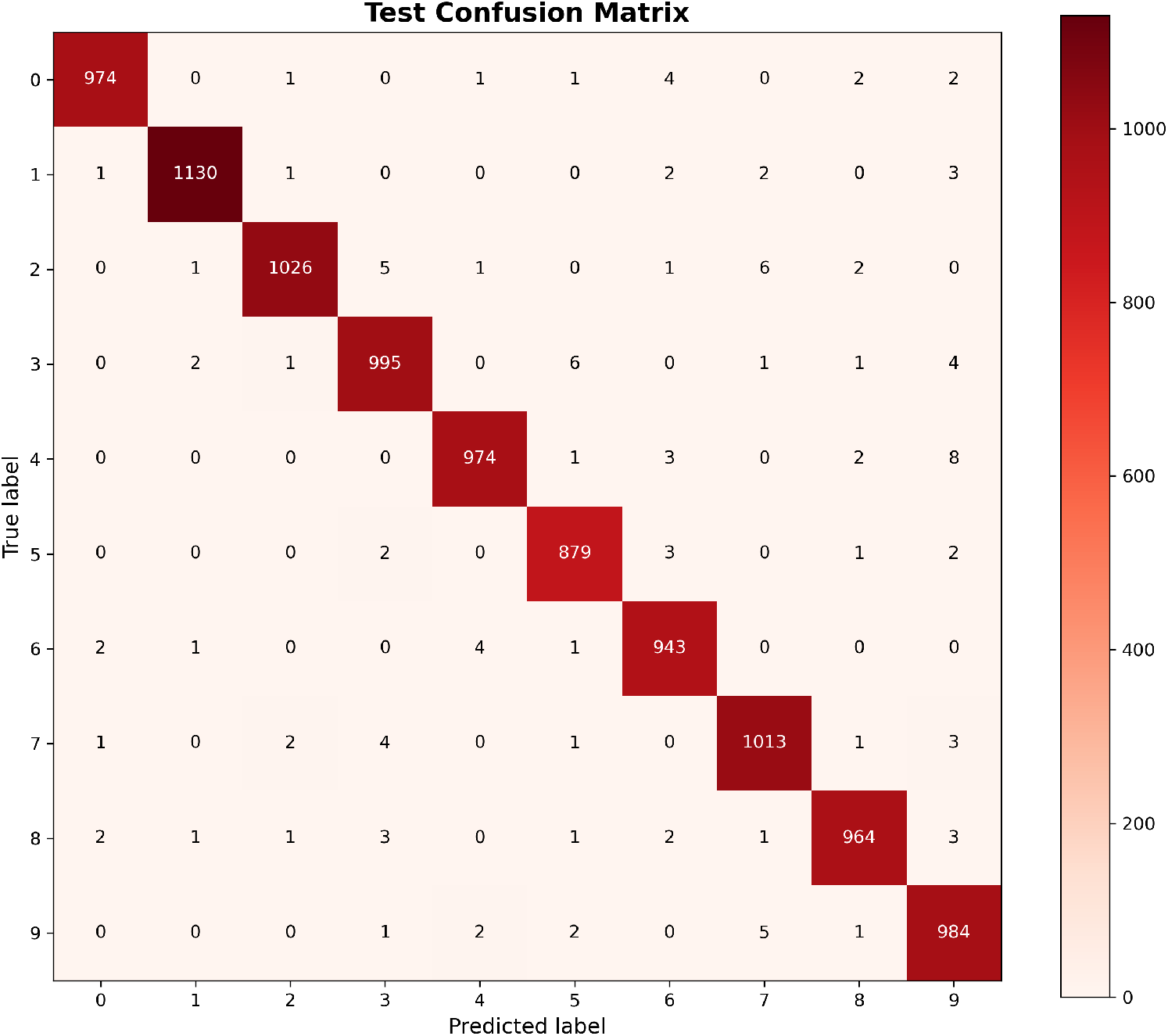
Test confusion matrix. Dark entries on the main diagonal count correctly classified samples, while all other entries count misclassifications

Beside a confusion matrix, one can build a bar plot of test mistakes per class to quickly visualise which classes cause the most mistakes (Figure 10). This graph can be a starting point for speculations on the network’s behaviour: for example, one might argue that the network makes most errors on samples from class “9” because those are more elaborate than samples from other classes, as they contain two different types of geometric primitives (a circle and a bar) that can come in different sizes or skews.

**Figure 10.**
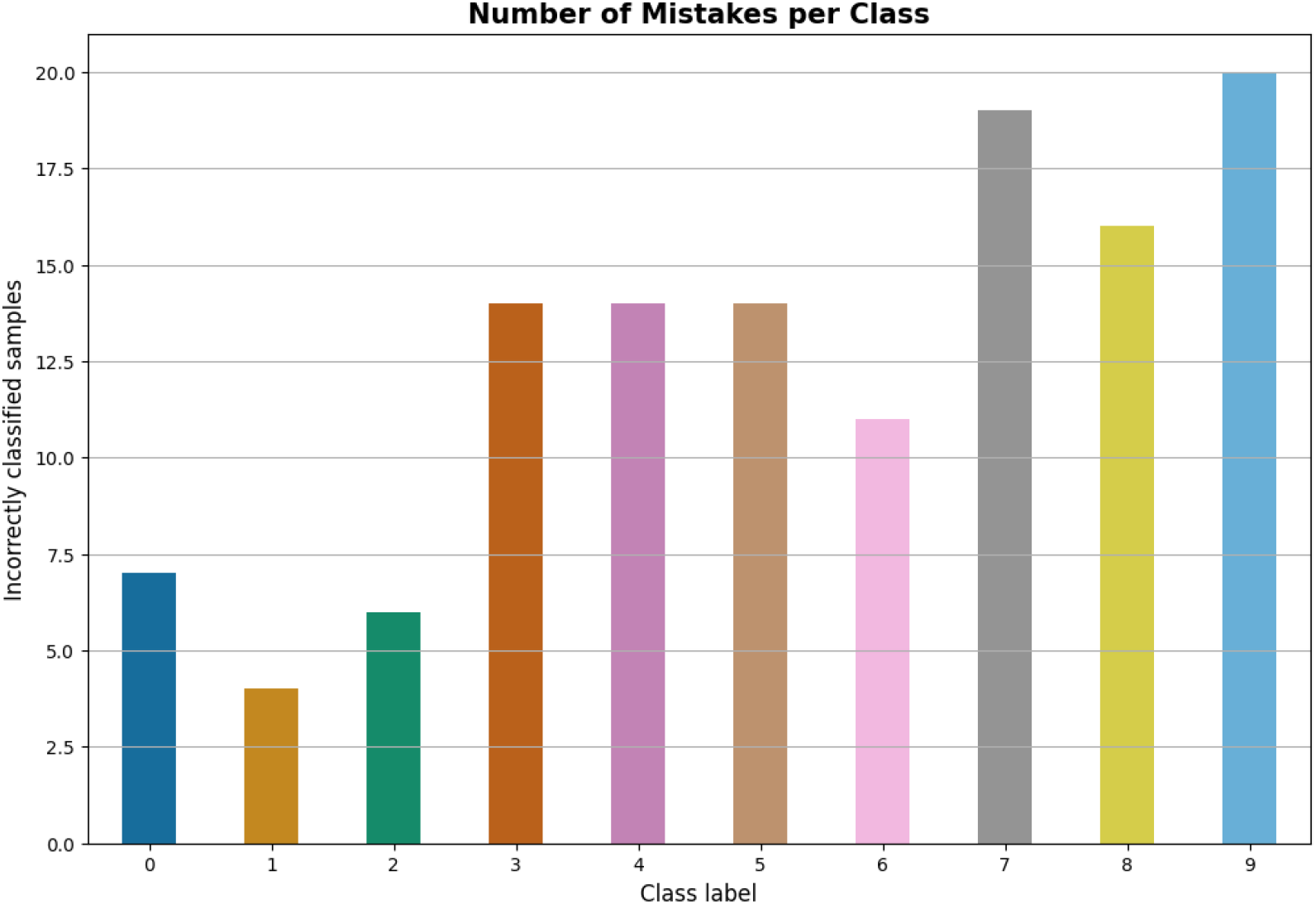
A bar plot of test mistakes per class

##### Code Example 19: The Test

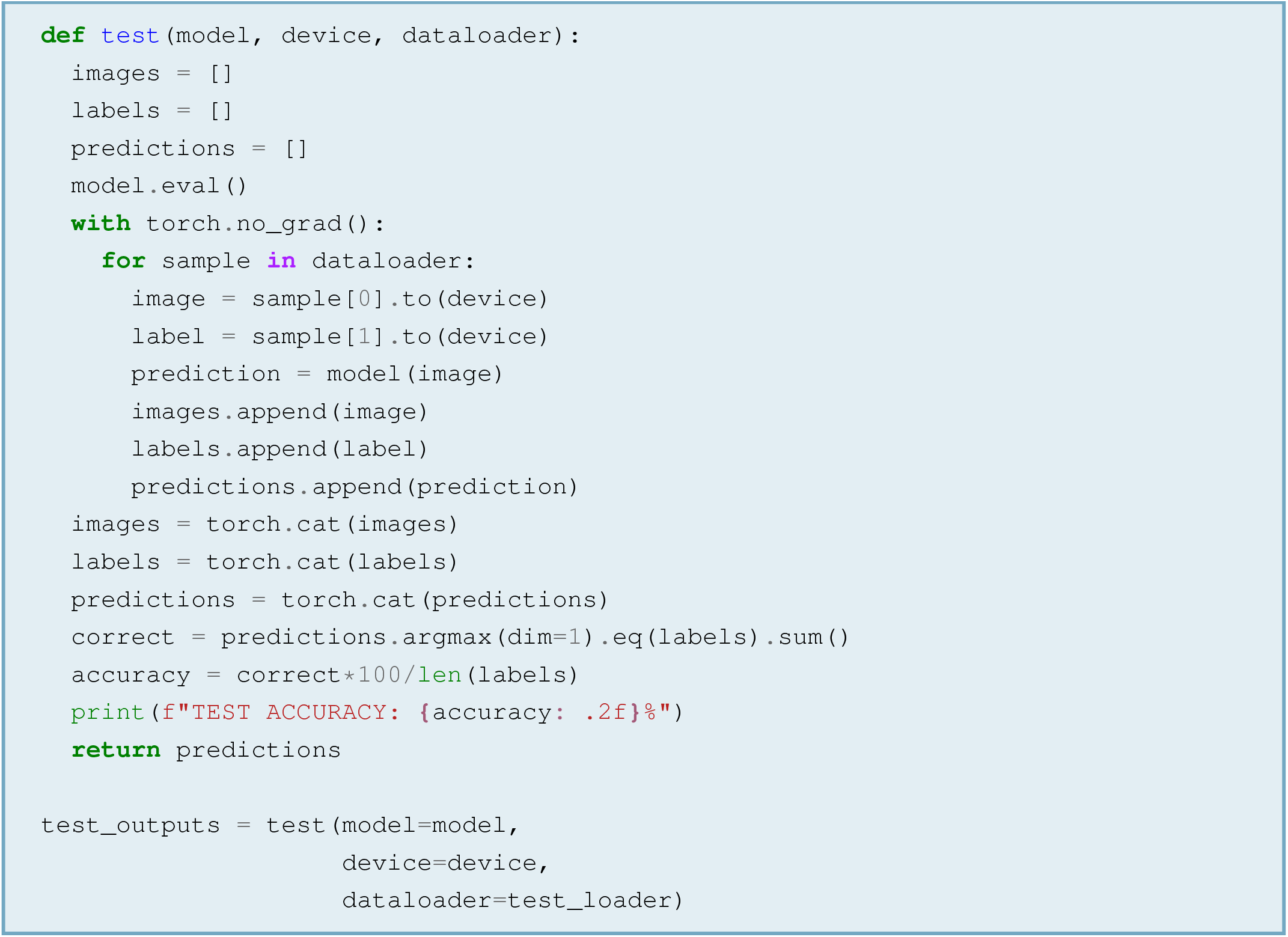

## 4 The Biological Plausibility of A/CNNs

As mentioned in previous sections, ANNs are mathematical models of networks of interacting biological neurons. Starting in the 1940s, they have evolved from objects of purely theoretical interest [1, 2, 41] to engineering tools that might have biological inspiration, but need not observe any biological constraints nor have neuroscientific relevance. Many innovations at the core of modern neural networks were born in this framework: for example, Yann LeCun and colleagues developed the first modern CNNs to recognise handwritten zip codes while working for the AT&T telecommunications company [42].

In the early 2010s, engineering-oriented CNNs started to produce human and superhuman results in complex object recognition tasks [5, 6]. Therefore, neuroscientists with a purely theoretical interest went back to studying neural networks after decades of indifference, adopting CNNs as models of biological vision and opening the debate about their biological plausibility — that is, whether (and to what extent) CNNs are able to represent the organisational and functional principles of biological brains. This section of the paper will analyse the plausibility of ANNs in general and CNNs in particular, covering every aspect from artificial neurons to energy consumption.

### 4.1 The Biological Plausibility of Artificial Neurons

Artificial neurons are highly simplistic models of the biological neuron, whose true complexity goes well beyond a simple weighted sum. Biological neurons have many morphological variants, each of which has peculiar functional characteristics [43]. Their structure is intricate, and the conventional division between dendrites, soma and axon is a mere approximation. The number and spatial distribution of dendrites vary greatly from one neuron to another, as do somatic topology and axon length. The configuration of dendrites impacts the receptive capacities of neurons, while the shape of the soma affects integration dynamics (that is, the way that post-synaptic potentials are integrated). Similarly, the speed of signal travel along axons depends on the axon’s length, section, detailed chemical composition, and degree of insulation, with myelination alone increasing speed by more than a hundred-fold [44]. Different neurons express different receptors and release different neurotransmitters, meaning that they exhibit different response profiles and different output effects. Finally, the number, type, and responsiveness of a neuron’s receptors may vary as a function of activity history, as may the number of dendrites and synapses. Instead of approximating a firing rate with a single scalar value, a detailed neuron model would take all those factors into account. However, a model like that would be completely useless. In fact, the high variability between neuron types and the ability of a neuron to change adaptively over time imply that highly detailed models could only represent a tiny portion of reality, with no hope of scaling up to the entire brain. Moreover, a perfectly detailed model would be a mere replication of the brain, so it would fail the primary goal of abstracting away its fundamental constituents.

Regardless of the thing it represents, a formal model must strike a balance between detail and abstraction: an excess of detail comes at the expense of generality, and excessive abstraction comes at the expense of realism [31]. This issue is particularly pressing for neuroscientific models, as the nervous system can be understood at many different scales — from molecules to brain-wide networks. Model requirements vary accordingly, with smaller scales requiring more detail and larger scales requiring more abstraction. This is due to detail being intrinsic to the smaller scales, where fine molecular dynamics are the very essence of phenomena. Such fine-grained details are often superfluous to the larger scales, whose essence is much coarser: for example, cognitive processes can be described very well without ever mentioning neurotransmitters. To bring clarity in this scenario, Herz et al. (2006) have proposed a taxonomy where neuroscientific models can belong to five different classes depending on the scale of interest [31]. The classes are called *levels* and are numbered from one to five, where one means *smaller scale* and five means *larger scale*. Models at smaller scales have a higher degree of detail, while models at larger scales have a higher degree of abstraction. However, all models are assumed to be realistic and provide an acceptable fit to empirical knowledge on nervous physiology.

- Level I models — or *detailed compartmental models* — aim to describe how neuronal ultrastructure gives rise to neuronal dynamics. To this end, they divide neurons into discrete compartments that are roughly homogeneous in structure and function and as such, can be studied in isolation at any level of detail. Level I models are powerful because they can approximate single-neuron dynamics with realism. However, their microscopic level of detail prevents any understanding of emergent phenomena like the formation of spike patterns. Moreover, their realism entails high computational complexity, preventing their use in simulations of large neuronal networks
- Level II models — also known as *reduced compartmental models* — follow the same logic as Level I models, but use one or very few compartments, assuming that neurons can be approximated as largely homogeneous. Compared to their *detailed* counterparts, *reduced* compartmental models can capture emergent phenomena like spike pattern generation and have lower computational complexity
- Level III models — also known as *single-compartment models* — ignore the neuron’s ultrastructure and focus entirely on understanding the generation of spikes from the interaction between ion currents. The Hodgkin-Huxley model of action potential generation is a single-compartment model, as it describes the action potential entirely as a function of sodium, potassium, and residual currents — with no regards for structural and functional differences across neuronal sectors
- Level IV models — or *cascade models* — describe neuronal activity as a cascade of events described by a sequence of mathematical functions, with no regards for the biological hardware that underpins those operations
- Level V models are also known as *black box models*. They describe neuronal activity in terms of conditional probabilities, meaning that they aim to estimate the probability that neurons produce a certain output spike train given that they received a certain input spike train. Relating one spike train to the other is useful for understanding the response profiles of neurons, and it has been instrumental in developing efficient coding theory [45]. However, it pays no attention to *how* an input is transformed into an output, treating the neuron as a complete black box

In this taxonomy, the artificial neuron of sits at Level IV. Indeed, it uses a linear operation (the dot product) to represent a spike train with a single number that approximates spike frequency, then it feeds that number into a non-linearity (for example, ReLU). This is enough to capture the capacity of biological neurons to receive an input with multiple components, weigh components differently, and distil them into a single output. The biological realism of the artificial neurons does not go any further. However, this is enough to construct models that can simulate a range of cognitive functions, from object recognition to complex decision making [5–16].

### 4.2 The Biological Plausibility of Artificial Neural Networks

Representing networks of neurons with the multiplication between a matrix of weights and a vector of data has a surprising degree of realism. As described at the beginning, the multiplication between a matrix and a vector is a collection of dot products between matrix rows and the single column of the vector. Despite concurring to the same result, those dot products are independent of each other — that is, they exert no mutual influences. In other words, the multiplication between a matrix and a vector is a parallel operation that is well-suited to the simulation of a parallel system like the brain.

The view of the brain as a parallel processor of multidimensional inputs is the hallmark of *parallel distributed processing (PDP)* [3, 4], a neurocognitive theory that construes cognition as an activation pattern over many neurons, which it models through matrix-vector multiplications. PDP emerged during the 1980s, posing itself in stark contrast with theoretical approaches that modelled cognition as the manipulation of abstract symbols or as series of discrete stages represented by box-and-arrow flowcharts [41]. Unlike other views of mental activity, PDP had two plausible traits. First, it grounded mental activity in the brain, conceptualising it as an emergent property of neuronal activity. Second, it took inspiration from the anatomical evidence in favour of parallel processing pathways across multiple brain systems. While very basic, these elements of plausibility have made the fortune of PDP models, transforming them into the spark that would light the artificial intelligence revolution.

The antithesis of PDP is *localist encoding*, where each neuron encodes a single input element. This means that in a localist brain, an *n*-dimensional input requires *n* neurons. If we take vision as an example and decide that the elements of visual space are fully-formed objects (and not photons), this means that the brain should have one neuron for all the possible variations of any single object, such as all the possible orientations of a body: indeed, a one-to-one correspondence between input elements and neurons implies that every neuron detects the presence of its target object, with no room for flexibility. Such a purely localist brain is implausible from the point of view of function as well as structure. From the point of view of function, the localist brain is incapable of recognising objects under different circumstances (which mammal brains can do), and from the point of view of structure, the localist brain should have either infinitely many neurons for infinitely many objects (which is impossible because the brain has finite size) or a finite number of neurons that increases at each new sensory experience (which is impossible because neurogenesis is not an instantaneous process). One may argue that human sensory space does not actually contain infinitely many objects, so a localist brain might be able to process all possible inputs. However, the statement that motion has infinite degrees of freedom cannot possibly be falsified, because we move in a continuous space. Thus, a localist motor system (where each neuron codes for a given muscle contraction strength) should indisputably have infinitely many elements, which is impossible for the reasons described above. Moreover, such a motor system would be incompatible with anatomical evidence that muscle contraction is controlled by mere hundreds of motor neurons, with contraction strength resulting from a distribution of activity over them [44].

An interesting trait of PDP models that separates them from the competition is their robustness to damage. When the representational power of an ANN is forcibly reduced (for example, through dropout), the network does not necessarily suffer a loss of function [38, 46]. This is a direct consequence of their parallelism: because all network units operate independently to achieve a common goal, non-ablated units can compensate the loss of the ablated ones. In contrast, ablating a step from a sequential model or a unit from a localist encoding makes the whole model useless. This fact contributes to making parallel distributed models more realistic than the alternatives, because human brain systems are resilient to damage and can adapt to partial loss or alteration of their parts [47].

In summary, artificial neural networks are PDP models of the brain. They are biologically plausible because they are based on the artificial neuron (which, in Herz et al.’s taxonomy, has Level-IV plausibility) and because they implement parallelism. Alternative models of brain function revolve around ideas like box-and-arrow flowcharts or localist encodings, which have less biological realism and have not translated into any useful implementations. However, ANNs/PDP models are unable to represent the connections between neurons within a given layer — which, in neural networks jargon, are known as *lateral connections*. This is certainly an unrealistic feature, but it does not compromise their ability to simulate cognitive functions with unparalleled success.

### 4.3 Hierarchical Processing: The Biological Plausibility of Deep Networks

In ANNs with more than one layer, a layer’s input is the next layer’s output, such that each layer receives more finely-processed data than the previous. This fact establishes a hierarchy of complexity, where the first layer processes the rawest data and the last one processes the most refined. This is similar to servants providing commodities to their masters in traditional social hierarchies, with wheat-harvesting farmers representing the input layer and bread-eating kings representing the output layer. As seen in section 2.1.4, such dynamics follow directly from the mathematics of matrix multiplication, which imply that each artificial neuron receives a weighted sum of all the activations in the previous layer. As a result, each neuron in a given layer pools the entire activity pattern of the previous layer, combining simpler patterns into more complex ones.

The hierarchical organisation of ANNs resonates with a wealth of anatomical, neurophysiological and neuropsychological findings that suggest the existence of hierarchical structures in the human brain. Such findings are particularly numerous in the visual system, which is a prototypical case study in neuroscience. At a very coarse level of analysis, the primate visual system has a two-tier structure:

- The first tier includes the eye, the lateral geniculate nuclei of the thalamus, the primary visual cortex, and other visual areas in the occipital lobes. This tier is anatomically compact, in the sense that each one of its elements is connected to the previous (either directly or indirectly)
- The second tier includes the ventral and dorsal visual streams, which are anatomically distinct and are thought to underpin two different visual functions [48]. The ventral stream (also known as *what pathway*) connects occipital visual areas to the temporal lobe and is thought to underpin visual object recognition — that is, the ability to associate semantic labels to visual objects. The dorsal stream (also known as *where* pathway) connects occipital visual areas to the parietal lobe and is thought to underpin the localisation of objects in space and sensorimotor transformations

Historically, the primary focus of vision neuroscience has been to understand visual object recognition — that is, the capacity to assign a semantic label to objects in visual space. For this reason, research has been focusing on the ventral more than the dorsal stream [49]. Here, studies across disciplines and levels of analysis have found convergent evidence in favour of a hierarchical organisation. The earliest evidence came from neurophysiological research on the domestic cat [50], whose primary visual cortex (V1) was shown to include neurons with a functional hierarchy: *simple cells* that responded preferentially to static oriented bars and *complex cells* that responded preferentially to moving oriented bars. Crucially, complex cells received input from a set of simple cells, possibly combining their static representations into a moving sequence. This ability to pool the inputs of several simple cells endowed complex cells with *translation invariance*, which is the ability to correctly perceive an object regardless of it moving. This finding inspired a wealth of conceptual and formal models, some of which laid the foundations for the current understanding of deep ANNs [51].

The idea of hierarchical organisation has been substantiated by findings that the macaque’s ventral stream has an anatomical hierarchy [52] and complex object perception emerges in later stages of ventral activity in both monkeys [53] and humans [54]. Moreover, lesion studies and neuropsychological experience suggest a functional dependence of later stages of the ventral stream on earlier ones: in monkeys like in humans, damage to earlier regions produces complete blindness to sectors of the visual field [55], while damage to later stages produces more specific, articulated deficits of complex object recognition [56–58].

Unfortunately, hierarchical models of vision have three important limitations:

- First, they do not consider the existence of recurrent and lateral connections in the brain (respectively, connections from higher to lower layers and within-layer connections). Nonetheless, the hierarchical model of the macaque visual system is implicitly *based* on the existence of recurrent connections, as it assigns a higher rank to structures that receive more forward connections than they send backward ones [59]. The very existence of those connections implies that they have a function, because unused synapses should be eliminated by standard metabolic processes [60]. Nonetheless, classic hierarchical models do not assign any role to recurrent connections, construing a purely feedforward stream of information. This is not a flaw *per se*, as it is natural for models to simplify reality. However, recurrent and lateral connections might be key to visual processing, and neglecting them might decrease the plausibility of ANNs to a degree that some could not accept. In particular, failing to incorporate recurrent connections might make ANNs unable to interpret ambiguous and complex natural scenes through the exploitation of previously acquired knowledge [59, 61]
- Second, hierarchical models assume that visual information follows a single route, always flowing forward along a given chain of invariant connections. However, the connections between biological neurons evolve continuously through plastic processes, modulating their functional properties. Moreover, visual pathways are densely connected with other systems of the brain, exchanging bidirectional influences that should continuously modulate structure and function [62]. Thus, modelling vision as a one-directional flow of information along an isolated stream may be overly simplistic
- Third, the empirical evidence in favour of hierarchical processing is less compelling than once thought. Hegdé and Van Essen (2007) [63] compared the response profiles of neurons in three different macaque visual areas that belonged to three different levels of the anatomical hierarchy (V1, V2, and V4). If the anatomical hierarchy had a clear functional correspondence, neurons in V1 would be selective to simpler stimuli than neurons in V2, which would in turn be tuned to simpler stimuli than neurons in V4. However, the authors found that all neurons responded to simple as well as complex shapes. This result suggests that visual processing is not necessarily hierarchical as ANNs would imply, and it paints a more complex picture than suggested by mainstream models

In summary, deep ANNs are biologically plausible because of their multi-layer, hierarchical organisation. However, they have important limitations related to the absence of recurrent and lateral connections, and some results suggest that hierarchy might not be as important as once thought for biological plausibility.

### 4.4 The Biological Plausibility of Convolutions: CNNs As Vision Models

Since their inception [42], deep CNNs have become a modelling standard in the computational cognitive neuroscience of vision, where convolution appears to be a powerful tool. Despite the simplicity of their basic structure, CNNs have elements of strong resemblance with biological vision — in particular, with its view as a hierarchical system [64].

In CNNs, each kernel is applied across a whole input sample, such that the same parameters are used to scan the entire input^15^. However, kernels project to a different neuron at each convolution step. This is somewhat plausible from the point of view of biology, as it simulates the fact that a finite quantity of sensory neurons represents an entire input space (akin to a fixed number of ganglion cells encoding information from the whole visual field). Moreover, the mathematics of convolution imply that artificial neurons use a single number (the activation) to represent a portion of the input as large as the kernel. This replicates the fact that each biological neuron represents its whole receptive field with a single signal, which is its firing rate.

After a convolutional layer, it is frequent for CNNs to apply a *pooling* operation, where the activations of neighbouring neurons are summarised (that is, *pooled*) into a single number (which is usually their maximum or their average). This operation introduces translation invariance in downstream neurons because it endows them with the ability to respond to a pattern regardless of its specific location within the receptive field. Moreover, it is an element of plausibility because it simulates the hierarchy found between simple and complex cells^16^.

Finally, CNNs are deep and hierarchically organised. This allows them to construct very abstract representations of the input, to the point of mapping complex multidimensional patterns onto single, pointwise responses like semantic labels produced by single output neurons. This can be seen as the progressive emergence of so-called *grandmother cells*, or neurons that fire selectively for a single input pattern [54]. As explained in the comparison between localist encoding and PDP models, the plausibility of grandmother cells is dubious. Nonetheless, the similarity between CNNs and hierarchical models of the ventral stream is striking, as are the performance capacities of CNNs. For this reason, it is worth investigating the nature of the representations learned by CNNs and how they compare to primate vision. This is a research frontier and there are few established facts. Perhaps the most solid finding is that the first layer of deep CNNs can learn Gabor-like features — that is, elongated regions of alternating luminance values that humans perceive as oriented bars. However, visualising and interpreting the representations learned by deeper layers is a non-trivial problem. One way to solve it is to use the *deconvolution method* [65], which inverts convolutions in order to find the image features that maximally drive response. Visualizations produced with the deconvolution method suggest that deeper neurons become selective to increasingly complex image features, but those results might be biased: in fact, they show which image features drive a unit’s response, but this does not mean that the unit is *selective* to those features — it only means that the unit is *able to represent them*. Moreover, units that represent interpretable features are often selected and perused because of their capacities, but they might well be an exception rather than the network’s standard.

One relatively robust finding is that CNNs construct similar representations to the ones of primate ventral streams [22, 64, 66]: for example, the representations learned by the AlexNet CNN [5] are similar to early occipital representations in the first few layers, and monotonically more similar to temporal representations across subsequent layers [67].

Despite their outstanding results, standard CNNs remain feedforward, hierarchical, and modality-specific models. As such, they fail to account for the existence of recurrent connections in the brain, as well as to capture the dynamical, varied, and multimodal nature of true nervous activity. They are the state of the art in the computational cognitive neuroscience of vision, but they leave much unexplained. One way to increase the explanatory power of A/CNNs is to introduce brain-inspired components in their structure. For example, recurrent convolutional neural networks (RCNNs) — that is, CNNs with feedback connections in their convolutional layers — have been shown to outperform standard, feedforward CNNs in object recognition tasks, emboldening the view that recurrence is required to explain vision [68, 69]. However, feedforward CNNs had already been shown to outperform humans in terms of visual recognition accuracy [6]. Therefore, RCNNs would only be closer to explaining biological vision if they were shown to achieve equal or better results than their feedforward counterparts, in spite of lower computational requirements or stronger plausibility constraints. However, studies that perform such detailed comparisons seem to be uncommon. Therefore, the only conclusion that can be drawn is that deep learning is a powerful modelling platform that allows scientists to test different views of brain function and compare results with relative ease. The evolution from traditional artificial neural networks, through feedforward CNNs and to RCNNs has demonstrated that the biological plausibility of deep learning can be progressively increased with the introduction of brain-inspired operations, thus suggesting that further steps in this direction could yield more realistic models of brain function.

### 4.5 The Biological Plausibility of Error-Driven Learning

Error backpropagation is the prototypical learning algorithm for artificial neural networks: while many more exist, this is certainly the most common. It has been analysed extensively during more than three decades of history, its mathematical treatment and code implementation are relatively simple, and its power has been shown by a wealth of successful models. However, it is widely believed to be biologically implausible.

A fundamental component of backpropagation is the computation of an error function that compares a model’s output with a target. Models are assigned a task (for example, classify an image as containing a dog or cat) and their performance is compared to an optimum (for example, assigning the correct label to the image). Subsequently, backpropagation is used to correct model parameters and nudge them towards the optimum.

Using backpropagation to model learning in the brain implies two strong assumptions:

- The first assumption is that the brain is organised into domain-specific systems — that is, sets of interacting neurons that carry out a single function like object recognition. In fact, backpropagation hones the performance of an A/CNN until it approximates perfection on a single task, as if there existed task-specific brain systems that learn to carry out their function independently from others. This contradicts anatomical evidence in favour of integration between brain regions, with all the neurons of the brain interacting with each other by varying degrees. It also contradicts the evidence that many cortical areas are multisensory, [62] and is therefore biologically implausible
- The second assumption is that cognition is goal-driven — that is, cognitive entities strive to approximate an exogenous target that exists independently of them. While there is no clear anatomical or physiological evidence to contradict it, this assumption is less than obvious. In fact, it proposes an *a priori* environment whose features are independent from the cognitive entity that perceives it and acts upon it, and to which the cognitive entity must adapt. Such proposed environment would have a neat structure with clear-cut perceptual and categorical boundaries, supporting strong dichotomies like correct vs. incorrect semantic labelling. Unfortunately, such net distinctions are less than pervasive in the real world, where sensory inputs can be assigned multiple semantic labels without mistake [70]. This is often due to biological cognitive agents bringing their previous knowledge and beliefs into perceptual judgements [71] — something that plain backpropagation does not take into account. In addition, the psychological literature and common experience are dense with non-linear phenomena like insight learning, which do not fit in the conceptual infrastructure of backpropagation. Therefore, cognition cannot be construed as merely stimulus reception followed by the production of a single, well-defined response (e.g., a semantic label). At most, this can be true of certain types of infant learning, where parents or other teachers instruct the infant about established facts of the environment where they were born. But even in this domain, the plausibility of backpropagation learning is restricted to the behavioural and cognitive levels: it does not extend to the neurophysiological level, where the assumption of segregated, task-specific brain systems remains invalid

Another reason why backpropagation is biologically implausible is that it imposes a dichotomy between *forward pass* (when a network processes data) and *backward pass* (when error derivatives are used to nudge network parameters in the optimal direction). The backward pass depends on the results from the forward pass, and the latter uses weights that have been adjusted by the backward pass. As a consequence, computations must be timed precisely to ensure a perfect alternation, where each pass builds on the results from the other [32]. In other words, once a neuron has been involved in the forward pass it should remain silent until involved in the backward pass, and then it should remain silent again until the right stage of the next forward pass. Such a precisely timed behaviour is not common in biological systems and if even it was, there would be no anatomical candidate to the role of master clock.

A third argument against backpropagation is that it requires derivatives of the error with respect to each weight. In a brain, this would require some implementation of weight parameters and a memory buffer to temporarily store weights and activations between the forward and backward passes. Conceptually, weight parameters model synaptic strengths, and their iterative modifications model the plastic processes that underpin biological learning. However, a synapse is a very complex system and plasticity is an extremely articulated process, which involves numerous cascades of chemical reactions at various timescales [44]. Therefore, there are no straightforward candidates for the role of weight parameters nor for the role of activation buffer.

A fourth argument is that backpropagation is based entirely on linear operations, but experimental evidence shows that nervous circuits implement numerous non-linearities [32]. Therefore, it is at least unlikely that learning could be entirely linear.

Finally, backpropagation learning is a resource-intensive process, which requires many data and many iterations of the input-output-weight correction cycle. Indeed, weight updates must be very small in order to avoid stability issues, so the path from the zero-knowledge start to task proficiency is walked in many steps. Moreover, models should be exposed to a rich and varied body of data, which some maintain should be ten times the parameters [22]. As large models can have millions of parameters, they should be exposed to tens of millions of input samples that they shall process over tens or hundreds of epochs. Therefore, a large model might easily be exposed to billions of input events during training. This is far from biological reality, where primates can learn to recognize an object after one brief presentation.

In summary, there are many reasons why backpropagation is biologically implausible. However, most of the arguments against it can be reasonably countered. First, neuroscience research is yet to find a mechanism by which single synaptic modifications coordinate to achieve an outcome at the network and behavioural levels [12]. Therefore, there is no evidence to falsify backpropagation as a network learning model, and the arguments against it are largely based on evidence from other scales or fields. For example, backpropagation’s assumption of a net divide between agent and environment is a matter of philosophy, with no direct relationship to network dynamics. Given the effectiveness of backpropagation in A/CNNs, it would be reckless to reject it on the grounds of a philosophical argument. In fact, backpropagation might well be valid as a partial theory of biological learning, whose extension or integration with other theories could provide a coherent framework that does not entail any problematic assumptions. This would be coherent with backpropagation-trained CNNs explaining certain aspects of cognition (e.g., core object recognition) but failing to explain others (such as one-shot learning or insight).

Similarly, it might be reckless to reject backpropagation on the grounds of no anatomical substrates. In fact, backpropagation is a learning rule at the network level, while the implementation of weights and activation buffers is a problem to solve at the single-cell level. In other words, it might be that learning by error feedback is an emergent property of networks that cannot be mapped precisely onto cellular structures or sub-cellular phenomena. The same holds for the objection that backpropagation is entirely linear, or that there is a clear alternation between forward and backward passes. In fact, it is not implausible that neurons take part in other tasks, remain at resting state, or oscillate towards an equilibrium between their involvement in the forward and backward passes.

In general, backpropagation must not be taken literally — that is, we should not expect to find a perfect fit with brain physiology. If we take it as the abstraction that it is, backpropagation has many elements of plausibility. The first one is the modification of synaptic strengths, whose importance for biological learning is established. The second is the use of feedback connections, which are pervasive in cortical anatomy. The role of such connections in network physiology is still unclear, so it is impossible to make strong statements about their influence on biological learning. However, the signals that they send are guaranteed to modulate spiking patterns in receiver neurons. In turn, these spiking patterns are guaranteed to influence synaptic strengths by means of plasticity. Thus, feedback connections are almost certain to play an instrumental role in biological learning [12]. A third element of plausibility is the exploitation of error signals, which reinforcement learning (Sutton & Barto, 1998) and successful computational neuroscience theories [72] propose as a biological learning primitive. Finally, backpropagation models have achieved good success at representing the response properties of neurons in the posterior parietal cortex [73], primary motor cortex [74], and ventral visual stream [67], making it hard to drop them.

### 4.6 Energy Consumption: The Biological Plausibility of Digital Computing

One last aspect to consider are the energy requirements of A/CNNs, which are largely different from those of human brains. The parallelism of matrix multiplications endows neural networks with great power, but requires just as much. Each dot product is a <u>weighted sum</u> that, depending on the size of the data and of neural network layers, can have tens or hundreds of terms. A single matrix-vector multiplication involves many such sums, meaning that every network layer requires hundreds or thousands multiplications and additions to be carried out in parallel. Many of those operations involve ones or zeros, so they take up computer memory and computation time in exchange for a small contribution to the final result. More memory and more time translate into more electric power, thus more energy requirements. This is a pressing issue for large deep networks, which include more processing layers and larger matrices compared to traditional networks. The iterative nature of learning and of scientific research makes requirements even larger, as a single model may undergo many thousand computing cycles before it reaches maturity. Strubell et al. (2019) [75] estimated that developing a single publication-level, state-of-the-art language model required them to go through almost five thousand versions of it, for a total expense in electricity of about ten-thousand dollars and an environmental footprint of more than 78000 pounds of carbon dioxide (that is, more than 35000 kilos). These numbers are considered representative of a full deep learning development cycle, and they are coherent with the finding that the compute demands of deep learning models have been doubling every 3.4 months between 2012 and 2018 — an increase in the hundreds of thousands over just six years (the first of the deep learning revolution) [76]. Set aside the unacceptably high carbon footprint, such results highlight a large discrepancy between the energy demands of biological brains and those of modern ANNs trained on digital computers.

The human brain is a metabolically demanding organ, but its absolute energy requirements are modest compared to deep learning’s. In the average adult human male, the brain amounts to only 2% of total body weight but consumes 20% of the body’s oxygen supply, as well as 20− 25% of total glucose. These inputs result in 0.25 kilocalories per minute, which correspond to about 20% of the body’s average metabolic rate. Thus, a single and relatively small organ takes up a fifth to a fourth of bodily energy sources and contributes for a fifth to the overall energy balance [77]. Luckily, the brain is much more efficient than ANNs: while the latter necessitate huge resources to perform a single function, the human brain sports full intelligence and coordinates an entire body on just a single meal. Nonetheless, concluding from these premises that deep learning systems have biologically implausible energy demands may be imprecise. In fact, their energy requirements may be disproportionate with respect to other software, but it is true that the energy requirements of the human brain are disproportionate with respect to those of other organs. Moreover, the comparison is biased by the structural differences between digital computers and the human brain, which have completely different hardware. While negligible in other contexts, this difference is key when it comes to energy. Comparing the mathematical descriptions of natural and artificial neurons is an abstract matter, as it is comparing natural and artificial learning algorithms: in both cases, the objects of comparison are not subject to any physical constraints, or models can be adapted to accommodate them. In contrast, energy use is an inherently physical aspect that cannot possibly be decoupled from hardware characteristics. Thus, the difference between the figures of natural and artificial systems may be due to computer hardware having lower energy efficiency than nervous tissue, and not to deep learning models being biologically implausible in concept. For this reason, important lines of research have been opened towards the goal of building new generations of computing hardware that more closely approximate the energetic properties of nervous tissue [78–81].

## 5 Lessons for the Neuroscience Community

The field of machine learning — of which A/CNNs are a small, but crucial part — has gone through a massive expansion over the last two decades, laying the grounds for the ongoing artificial intelligence (AI) revolution. Among the many factors that catalysed this explosion, the following are certainly the most important:

- Commercial, social, and scientific interest in the technology, which attracted large investments from companies and, to a minor extent, universities and other institutions
- The availability of large-scale data from the Internet (chiefly images, text, and video)
- The availability of relatively cheap parallel computing hardware (chiefly CUDA-enabled GPUs by NVIDIA)
- The availability of high-quality free and open source software (like PyTorch and Tensorflow)
- The wide adoption of agile, fast, and free (or low-cost) publication strategies (like arXiv preprints or conference proceedings)

With the convergence of these factors, neural networks have gone from a relatively uncommon theory and limited engineering tools, to gold-standard vision models and actors of the AI revolution. Their story shows that investing in technology, sharing data, sharing methods, adopting agile publication practices and paying researchers well can generate outstanding progress. Importantly, this can happen regardless of previous expectations: until the AlexNet CNN [5] triumphed at the ImageNet Large Scale Visual Recognition Challenge in 2012, very few people (if anyone) suspected that neural networks would become pervasive within the following ten years.

Like neural networks in the late twentieth and early twenty-first century, neuroscience is a fascinating and potentially impactful field. However, it struggles with methodological issues (like non-reproducible research practices), an insufficiently *open* culture, and a lack of funding that drives researchers away from the field or forces them to work on the problems that they can afford (rather than the problems that matter). Hopefully, neuroscientists can learn from neural networks and work towards better progress: searching impactful solutions for big problems, working reproducibly, sharing data, sharing reproducible code, and pushing towards sustainable publication practices. More investments might follow.

## Acknowledgements

This paper is based on material that Matteo De Matola (MDM) prepared for and taught in Lecture 3.3 of *Python for (open) Neuroscience*: a course by Luigi Petrucco at the University of Trento’s Doctoral School in Cognitive & Brain Sciences (Spring 2024). Thanks to Dr. Petrucco for the chance of teaching that lecture: without it, this paper would not exist.

The paper’s title is partly a tribute to Jackson & Bolger’s 2014 EEG tutorial [82]. While their work is completely unrelated to A/CNNs, the two have made a great job at disseminating the physics and physiology of EEG to a broad audience, inspiring some of this tutorial’s style.

Section 4 (*The Biological Plausibility of A/CNNs*) is adapted from MDM’s master’s thesis, [A study of spiking neural networks for biologically plausible deep learning](https://thesis.unipd.it/handle/20.500.12608/3104), defended at the University of Padova on 25 November 2021.

Thanks to prof. Carlo Miniussi (University of Trento) for reading an early version of this manuscript and providing constructive feedback.

Finally, thanks to everyone who has ever published freely available educational materials: this paper is a humble attempt at giving something back.

## Code Availability

This paper has a companion Jupyter Notebook that can be found on GitHub at this link. The notebook includes all code examples and the associated textual explanations, organised in lecture format.

## Conflict of Interest

The authors declare no conflicts of interest.

## Declaration of Authorship

This work is entirely the product of the authors’ own initiative, intellect, effort, and time, and the authors are entirely responsible for its contents. No artificial intelligence tools were used at any stage of preparation.

## Footnotes

1 As revealed by searching *artificial neural network AND (neuroscience OR neural OR brain OR cortex OR cognitive)* on PubMed in January 2026 and constraining results to the last five years

2 A Jupyter Notebook is a document that alternates blocks of static text to blocks of executable Python code. It can either be downloaded and used locally (installations required) or opened and run in a web service like [Google Colab](https://colab.research.google.com/) (no installations required)

3 See the web companion for a definition of these terms. However, note that understanding them is irrelevant at this stage

4 A weighted sum is a sum of elements that are multiplied by a number (called *weight*) before being added together

5 [NumPy](https://numpy.org/) is a Python library for vector and matrix operations

6 The key to seeing these facts is that 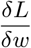 is multiplied by negative *α* and the product of two negative quantities is a positive, while the product of a negative and a positive quantity is a negative

7 There are many different flavours of backpropagation, each one using a slightly (but nonetheless, significantly) different formulation of the weight-updating rule

8 *k* = 10 and *k* = 5 seem to work well empirically. Using *k* = *N*, where N is the total number of observations, is referred to as *leave-one-out cross-validation (LOOCV)* because only one item is left out from the training at every cross-validation iteration

9 Cross-validation can be very costly from a computational perspective: for each of *k* partitions, a model must do training and validation on a dataset that includes hundreds or thousands of samples — and this is repeated for multiple candidate models in the case of model selection

10 Differentiation is the act of taking the <u>derivative</u> of a function. How tensors — which, in essence, are just containers of numbers — can represent a differentiable function is beyond the scope of this paper, whose goal is to demystify the *science* behind A/CNNs more than the engineering. Interested readers can refer to the [official PyTorch documentation](https://docs.pytorch.org/tutorials/beginner/basics/autogradqs_tutorial.html) and to [this](https://colah.github.io/posts/2015-08-Backprop/) excellent article by Anthropic researcher Chris Olah

11 A deep understanding of this code example requires to know Python classes, which are beyond the scope of this paper. Interested readers can refer to introductory materials like [this Real Python article](https://web.archive.org/web/20260131092830/https://realpython.com/python-classes/). Others can just read through the code example and rely on the docstrings — that is, the explanations in triple quotes (“““)

12 This is done by a softmax function, which in this case is implemented within nn.CrossEntropyLoss. Other implementations might require to apply the softmax explicitly and subsequently feed the result through a cross-entropy function. Note that nn.CrossEntropyLoss does not exactly use a softmax but a log-softmax function, which is merely the logarithm of the softmax. Using a log-softmax makes sense because it is more computationally efficient and because a logarithm would be involved in the subsequent calculation of cross-entropy anyway

13 In PyTorch, all optimisers reside in the ‘optim’ package.

14 In this case, the CNN was regularised with dropout and weight decay. Weight decay is a part of Adam’s weight-update equation, while dropout is built into the network as a layer. Another example of a built-in regularisation strategy is *batch normalisation*, which is not used in this tutorial. Setting the network into training mode enables any built-in regularisation strategies, while setting it into validation mode disables them

15 This is known as *parameter sharing* in the deep learning literature

16 Note that pooling operations can be replaced by convolution strides larger than one. Therefore, translation invariance can be achieved without pooling if stride is set to 2 or more

## Notes

https://github.com/matteo-d-m/cnns-tutorial

### Competing Interest Statement

The authors have declared no competing interest.

## References

[1] Warren S McCulloch and Walter Pitts. A logical calculus of the ideas immanent in nervous activity. The Bulletin of Mathematical Biophysics, 5(4):115–133, 1943.

[2] Marvin Minsky and Seymour Papert. Perceptron: An introduction to computational geometry. The MIT Press, Cambridge, Expanded Edition, 19(88):2, 1969.

[3] David E Rumelhart, James L McClelland, PDP Research Group, et al. Parallel distributed processing, Volume 1. Explorations in the Microstructure of Cognition: Foundations. MIT Press, 1986.

[4] James L McClelland, David E Rumelhart, PDP Research Group, et al. Parallel distributed processing, Volume 2. Explorations in the Microstructure of Cognition: Psychological and Biological Models, volume 2. MIT Press, 1987.

[5] Alex Krizhevsky, Ilya Sutskever, and Geoffrey E Hinton. Imagenet classification with deep convolutional neural networks. Advances in Neural Information Processing Systems, 25, 2012.

[6] Kaiming He, Xiangyu Zhang, Shaoqing Ren, and Jian Sun. Deep residual learning for image recognition. In Proceedings of the IEEE Conference on Computer Vision and Pattern Recognition, pages 770–778, 2016.

[7] Gao Huang, Zhuang Liu, Laurens Van Der Maaten, and Kilian Q Weinberger. Densely connected convolutional networks. In Proceedings of the IEEE Conference on Computer Vision and Pattern Recognition, pages 4700–4708, 2017.

[8] Jeffrey L Elman. Finding structure in time. Cognitive Science, 14(2):179–211, 1990.

[9] Sepp Hochreiter and Jürgen Schmidhuber. Long short-term memory. Neural Computation, 9(8):1735–1780, 1997.

[10] Ashish Vaswani, Noam Shazeer, Niki Parmar, Jakob Uszkoreit, Llion Jones, Aidan N Gomez, Łukasz Kaiser, and Illia Polosukhin. Attention is all you need. Advances in Neural Information Processing Systems, 30, 2017.

[11] Tom Brown, Benjamin Mann, Nick Ryder, Melanie Subbiah, Jared D Kaplan, Prafulla Dhariwal, Arvind Neelakantan, Pranav Shyam, Girish Sastry, Amanda Askell, et al. Language models are few-shot learners. Advances in neural information processing systems, 33:1877–1901, 2020.

[12] Timothy P Lillicrap, Adam Santoro, Luke Marris, Colin J Akerman, and Geoffrey Hinton. Backpropagation and the brain. Nature Reviews Neuroscience, 21(6):335–346, 2020.

[13] Volodymyr Mnih, Koray Kavukcuoglu, David Silver, Andrei A Rusu, Joel Veness, Marc G Bellemare, Alex Graves, Martin Riedmiller, Andreas K Fidjeland, Georg Ostrovski, et al. Human-level control through deep reinforcement learning. Nature, 518(7540):529–533, 2015.

[14] Timothy P Lillicrap, Jonathan J Hunt, Alexander Pritzel, Nicolas Heess, Tom Erez, Yuval Tassa, David Silver, and Daan Wierstra. Continuous control with deep reinforcement learning. arXiv preprint arXiv:1509.02971, 2015.

[15] David Silver, Aja Huang, Chris J Maddison, Arthur Guez, Laurent Sifre, George Van Den Driessche, Julian Schrittwieser, Ioannis Antonoglou, Veda Panneershelvam, Marc Lanctot, et al. Mastering the game of go with deep neural networks and tree search. Nature, 529(7587):484–489, 2016.

[16] David Silver, Julian Schrittwieser, Karen Simonyan, Ioannis Antonoglou, Aja Huang, Arthur Guez, Thomas Hubert, Lucas Baker, Matthew Lai, Adrian Bolton, et al. Mastering the game of go without human knowledge. Nature, 550(7676):354–359, 2017.

[17] Adam Paszke, Sam Gross, Francisco Massa, Adam Lerer, James Bradbury, Gregory Chanan, Trevor Killeen, Zeming Lin, Natalia Gimelshein, Luca Antiga, et al. Pytorch: An imperative style, highperformance deep learning library. Advances in Neural Information Processing Systems, 32, 2019.

[18] Mark S Goldman and Michale S Fee. Computational training for the next generation of neuroscientists. Current Opinion in Neurobiology, 46:25–30, 2017.

[19] Ashley Juavinett. Learning how to code while analyzing an open access electrophysiology dataset. Journal of Undergraduate Neuroscience Education, 19(1):A94, 2020.

[20] Ashley L Juavinett. The next generation of neuroscientists needs to learn how to code, and we need new ways to teach them. Neuron, 110(4):576–578, 2022.

[21] Ashley L Juavinett. Integrating programming into neuroscience courses. Journal of Undergraduate Neuroscience Education, 22(2):A99, 2024.

[22] Daniel LK Yamins and James J DiCarlo. Using goal-driven deep learning models to understand sensory cortex. Nature Neuroscience, 19(3):356–365, 2016.

[23] Fabian Isensee, Paul F Jaeger, Simon AA Kohl, Jens Petersen, and Klaus H Maier-Hein. nnu-net: a self-configuring method for deep learning-based biomedical image segmentation. Nature Methods, 18(2):203–211, 2021.

[24] Subathra Gunasekaran, Prabin Selvestar Mercy Bai, Sandeep Kumar Mathivanan, Hariharan Rajadurai, Basu Dev Shivahare, and Mohd Asif Shah. Automated brain tumor diagnostics: Empowering neuro-oncology with deep learning-based mri image analysis. Plos One, 19(8):e0306493, 2024.

[25] Yuki Wong, Eileen Lee Ming Su, Che Fai Yeong, William Holderbaum, and Chenguang Yang. Brain tumor classification using mri images and deep learning techniques. PloS One, 20(5):e0322624, 2025.

[26] Robin Tibor Schirrmeister, Jost Tobias Springenberg, Lukas Dominique Josef Fiederer, Martin Glasstetter, Katharina Eggensperger, Michael Tangermann, Frank Hutter, Wolfram Burgard, and Tonio Ball. Deep learning with convolutional neural networks for eeg decoding and visualization. Human Brain Mapping, 38(11):5391–5420, 2017.

[27] Alexander Craik, Yongtian He, and Jose L Contreras-Vidal. Deep learning for electroencephalogram (eeg) classification tasks: a review. Journal of Neural Engineering, 16(3):031001, 2019.

[28] Yannick Roy, Hubert Banville, Isabela Albuquerque, Alexandre Gramfort, Tiago H Falk, and Jocelyn Faubert. Deep learning-based electroencephalography analysis: a systematic review. Journal of Neural Engineering, 16(5):051001, 2019.

[29] Lukas AW Gemein, Robin T Schirrmeister, Patryk Chrabaszcz, Daniel Wilson, Joschka Boedecker, Andreas Schulze-Bonhage, Frank Hutter, and Tonio Ball. Machine-learning-based diagnostics of eeg pathology. NeuroImage, 220:117021, 2020.

[30] Charles R Harris, K Jarrod Millman, Stéfan J Van Der Walt, Ralf Gommers, Pauli Virtanen, David Cournapeau, Eric Wieser, Julian Taylor, Sebastian Berg, Nathaniel J Smith, et al. Array programming with numpy. Nature, 585(7825):357–362, 2020.

[31] A.V. Herz, T. Gollisch, C.K. Machens, and D. Jaeger. Modeling single-neuron dynamics and computations: a balance of detail and abstraction. Science, 314(5796):80–85, 2006.

[32] Yoshua Bengio, Dong-Hyun Lee, Jorg Bornschein, Thomas Mesnard, and Zhouhan Lin. Towards biologically plausible deep learning. arXiv preprint arXiv:1502.04156, 2015.

[33] Gareth James, Daniela Witten, Trevor Hastie, and Robert Tibshirani. An introduction to statistical learning: with applications in R. Springer, 2013.

[34] Martín Abadi, Ashish Agarwal, Paul Barham, Eugene Brevdo, Zhifeng Chen, Craig Citro, Greg S Corrado, Andy Davis, Jeffrey Dean, Matthieu Devin, et al. Tensorflow: Large-scale machine learning on heterogeneous distributed systems. arXiv preprint arXiv:1603.04467, 2016.

[35] Matthias Fey and Jan Eric Lenssen. Fast graph representation learning with pytorch geometric. arXiv preprint arXiv:1903.02428, 2019.

[36] Yann LeCun. The mnist database of handwritten digits. http://yann.lecun.com/exdb/mnist/, 1998.

[37] Ian Goodfellow, Yoshua Bengio, and Aaron Courville. Deep Learning. MIT Press, 2016. http://www.deeplearningbook.org.

[38] Nitish Srivastava, Geoffrey Hinton, Alex Krizhevsky, Ilya Sutskever, and Ruslan Salakhutdinov. Dropout: a simple way to prevent neural networks from overfitting. The Journal of Machine Learning Research, 15(1):1929–1958, 2014.

[39] Takuya Akiba, Shotaro Sano, Toshihiko Yanase, Takeru Ohta, and Masanori Koyama. Optuna: A next-generation hyperparameter optimization framework. In Proceedings of the 25th ACM SIGKDD International Conference on Knowledge Discovery & Data Mining, pages 2623–2631, 2019.

[40] Diederik P Kingma and Jimmy Ba. Adam: A method for stochastic optimization. arXiv preprint arXiv:1412.6980, 2014.

[41] James L McClelland. Connectionist models and psychological evidence. Journal of memory and language, 27(2):107–123, 1988.

[42] Yann LeCun, Bernhard Boser, John S Denker, Donnie Henderson, Richard E Howard, Wayne Hubbard, and Lawrence D Jackel. Backpropagation applied to handwritten zip code recognition. Neural Computation, 1(4):541–551, 1989.

[43] Rodolfo R Llinás. The intrinsic electrophysiological properties of mammalian neurons: insights into central nervous system function. Science, 242(4886):1654–1664, 1988.

[44] Eric R. Kandel, James H. Schwartz, Thomas M. Jessell, Steven Siegelbaum, and A. James Hudspeth. Principles of Neural Science. McGraw-Hill, New York, 5th edition, 2012.

[45] Horace Barlow. Redundancy reduction revisited. Network: Computation in Neural Systems, 12(3):241, 2001.

[46] Song Han, Huizi Mao, and William J Dally. Deep compression: Compressing deep neural networks with pruning, trained quantization and huffman coding. arXiv preprint arXiv:1510.00149, 2015.

[47] Simona Fiori and Andrea Guzzetta. Plasticity following early-life brain injury: Insights from quantitative mri. In Seminars in Perinatology, volume 39, pages 141–146. Elsevier, 2015.

[48] Melvyn A Goodale and A David Milner. Separate visual pathways for perception and action. Trends in Neurosciences, 15(1):20–25, 1992.

[49] James J DiCarlo, Davide Zoccolan, and Nicole C Rust. How does the brain solve visual object recognition? Neuron, 73(3):415–434, 2012.

[50] David H Hubel and Torsten N Wiesel. Receptive fields, binocular interaction and functional architecture in the cat’s visual cortex. The Journal of Physiology, 160(1):106, 1962.

[51] Maximilian Riesenhuber and Tomaso Poggio. Hierarchical models of object recognition in cortex. Nature Neuroscience, 2(11):1019–1025, 1999.

[52] D.J. Felleman and D.C. Van Essen. Distributed hierarchical processing in the primate cerebral cortex. Cerebral Cortex, 1(1):1–47, 1991.

[53] Maxime Cauchoix, Sébastien M Crouzet, Denis Fize, and Thomas Serre. Fast ventral stream neural activity enables rapid visual categorization. NeuroImage, 125:280–290, 2016.

[54] R Quian Quiroga, Leila Reddy, Gabriel Kreiman, Christof Koch, and Itzhak Fried. Invariant visual representation by single neurons in the human brain. Nature, 435(7045):1102–1107, 2005.

[55] Petra Stoerig and Alan Cowey. Blindsight in man and monkey. Brain: A Journal of Neurology, 120(3):535–559, 1997.

[56] Peter H Schiller. Effect of lesions in visual cortical area v4 on the recognition of transformed objects. Nature, 376(6538):342–344, 1995.

[57] Martha J Farah. Visual agnosia: disorders of object recognition and what they tell us about normal vision. The MIT Press, 1990.

[58] David Pitcher, Lucie Charles, Joseph T Devlin, Vincent Walsh, and Bradley Duchaine. Triple dissociation of faces, bodies, and objects in extrastriate cortex. Current Biology, 19(4):319–324, 2009.

[59] Jay Hegde and Daniel J Felleman. Reappraising the functional implications of the primate visual anatomical hierarchy. The Neuroscientist, 13(5):416–421, 2007.

[60] Won-Suk Chung, Nicola J Allen, and Cagla Eroglu. Astrocytes control synapse formation, function, and elimination. Cold Spring Harbor perspectives in biology, 7(9):a020370, 2015.

[61] Alberto Testolin and Marco Zorzi. Probabilistic models and generative neural networks: Towards an unified framework for modeling normal and impaired neurocognitive functions. Frontiers in Computational Neuroscience, 10:73, 2016.

[62] Asif A Ghazanfar and Charles E Schroeder. Is neocortex essentially multisensory? Trends in Cognitive Sciences, 10(6):278–285, 2006.

[63] Jay Hegdé and David C Van Essen. A comparative study of shape representation in macaque visual areas v2 and v4. Cerebral Cortex, 17(5):1100–1116, 2007.

[64] Nikolaus Kriegeskorte. Deep neural networks: a new framework for modeling biological vision and brain information processing. Annual Review of Vision Science, 1(1):417–446, 2015.

[65] Matthew D Zeiler and Rob Fergus. Visualizing and understanding convolutional networks. In European Conference on Computer Vision, pages 818–833. Springer, 2014.

[66] Daniel L Yamins, Ha Hong, Charles Cadieu, and James J DiCarlo. Hierarchical modular optimization of convolutional networks achieves representations similar to macaque it and human ventral stream. Advances in Neural Information Processing Systems, 26, 2013.

[67] Seyed-Mahdi Khaligh-Razavi and Nikolaus Kriegeskorte. Deep supervised, but not unsupervised, models may explain it cortical representation. PLoS Computational Biology, 10(11):e1003915, 2014.

[68] Ming Liang and Xiaolin Hu. Recurrent convolutional neural network for object recognition. In Proceedings of the IEEE Conference on Computer Vision and Pattern Recognition, pages 3367–3375, 2015.

[69] Courtney J Spoerer, Patrick McClure, and Nikolaus Kriegeskorte. Recurrent convolutional neural networks: a better model of biological object recognition. Frontiers in Psychology, 8:1551, 2017.

[70] Edwin G Boring. A new ambiguous figure. The American Journal of Psychology, 1930.

[71] Michael ER Nicholls, Owen Churches, and Tobias Loetscher. Perception of an ambiguous figure is affected by own-age social biases. Scientific Reports, 8(1):12661, 2018.

[72] Daniel M Wolpert and Zoubin Ghahramani. Computational principles of movement neuroscience. Nature Neuroscience, 3(11):1212–1217, 2000.

[73] David Zipser and Richard A Andersen. A back-propagation programmed network that simulates response properties of a subset of posterior parietal neurons. Nature, 331(6158):679–684, 1988.

[74] Timothy P Lillicrap and Stephen H Scott. Preference distributions of primary motor cortex neurons reflect control solutions optimized for limb biomechanics. Neuron, 77(1):168–179, 2013.

[75] Emma Strubell, Ananya Ganesh, and Andrew McCallum. Energy and policy considerations for deep learning in nlp. In Proceedings of the 57th Annual Meeting of the Association for Computational Linguistics, pages 3645–3650, 2019.

[76] Roy Schwartz, Jesse Dodge, Noah A Smith, and Oren Etzioni. Green ai. Communications of the ACM, 63(12):54–63, 2020.

[77] S. A. Huettel, A. W. Song, and G. McCarthy. Functional Magnetic Resonance Imaging. Sinauer Associates, Sunderland, MA, 2014.

[78] Giacomo Indiveri, Bernabé Linares-Barranco, Tara Julia Hamilton, André van Schaik, Ralph Etienne-Cummings, Tobi Delbruck, Shih-Chii Liu, Piotr Dudek, Philipp Häfliger, Sylvie Renaud, et al. Neuromorphic silicon neuron circuits. Frontiers in Neuroscience, 5:73, 2011.

[79] Yichen Shen, Nicholas C Harris, Scott Skirlo, Mihika Prabhu, Tom Baehr-Jones, Michael Hochberg, Xin Sun, Shijie Zhao, Hugo Larochelle, Dirk Englund, et al. Deep learning with coherent nanophotonic circuits. Nature Photonics, 11(7):441–446, 2017.

[80] Chetan Singh Thakur, Jamal Lottier Molin, Gert Cauwenberghs, Giacomo Indiveri, Kundan Kumar, Ning Qiao, Johannes Schemmel, Runchun Wang, Elisabetta Chicca, Jennifer Olson Hasler, et al. Largescale neuromorphic spiking array processors: A quest to mimic the brain. Frontiers in Neuroscience, 12:891, 2018.

[81] Mohammed A Zidan, John Paul Strachan, and Wei D Lu. The future of electronics based on memristive systems. Nature Electronics, 1(1):22–29, 2018.

[82] Alice F Jackson and Donald J Bolger. The neurophysiological bases of eeg and eeg measurement: A review for the rest of us. Psychophysiology, 51(11):1061–1071, 2014.

